# Global propagation of genetic perturbation effects through genome-wide stoichiometry conservation architecture

**DOI:** 10.1101/2025.05.26.656078

**Authors:** Genta Chiba, Ken-ichiro F. Kamei, Arisa H. Oda, Kunihiro Ohta, Yuichi Wakamoto

**Author notes:** These authors contributed equally to this work (Genta Chiba, Ken-ichiro F. Kamei).

## Abstract

Understanding how specific genetic changes lead organisms to different phenotypes is the cornerstone of genetics. However, even a single gene perturbation can cause global changes in gene expression profiles. Consequently, predicting which genes will be affected by a particular genetic perturbation remains a challenge. Here, we analyze changes in genome-wide gene expression profiles caused by single-gene perturbations in bacterial, worm, and human cells and show a simple rule that genes that intrinsically maintain the ratio of expression levels (stoichiometry) with a perturbed gene are more likely to be affected when it is deleted or inactivated. Furthermore, deleting a gene that maintains their stoichiometry with more genes tend to affect the expression levels of other genes genome-wide. Stoichiometry-conserving gene clusters can be found beyond local transcriptional regulatory units and metabolic pathways. Thus, the effects of genetic perturbations are propagated to other genes through a multi-level stoichiometry-conserving architecture in cells.

## Introduction

Cells respond to the changes in extracellular and intracellular conditions through coordinated modulation of their molecular composition to maintain their function and adapt (*1–5*). Genetic perturbations, such as point mutations and the deletion of a specific gene, can alter gene expression profiles and generate the phenotypic diversity that drives the adaptive evolution of cells and organisms. Typically, the effects of local genetic perturbations are not limited to the perturbed genes or their neighbors; even when a single gene is perturbed, its effects may propagate to many other genes and pathways, complicating genotype-phenotype correspondence (*6–9*). The complexity of global changes in gene expression profiles stems from the fact that genes in cells work cooperatively and interact with many other genes. Nonlinear relationships such as pleiotropy and epistasis are common in gene function relationships; local functional and regulatory relationships of a perturbed gene are usually insufficient to explain global transcriptional changes caused by the perturbation (*10–12*).

Systematic analyses of changes in cellular phenotype due to genetic perturbations have identified several features of the underlying transcriptomic and metabolic changes (*2*, *6*, *8*, *13*, *14*). For example, Perturb-seq, a method that combines multiplexed CRISPR-mediated gene inactivation and subsequent single-cell RNA sequencing (scRNA-seq), enables high-throughput assessment of the comprehensive gene expression phenotype for each genetic perturbation and has characterized genome-wide changes in gene expression profiles caused by single-gene inactivation (*13*, *15*, *16*). Importantly, the global changes in gene expression profiles are not random; the studies suggest the role of genetic architectures, such as gene modules, co-expression, and common regulation in modulating global gene expression profiles (*17*, *18*). Previous research on, for example, *Saccharomyces cerevisiae* and *Escherichia coli* has characterized novel functional genes by examining the transcriptional and metabolic profile similarities between genes involved in functionally related processes (*7*, *8*, *19*, *20*). However, despite these efforts, the system-level principle that explains how single-gene perturbations lead to specific genome-wide changes in gene expression still remains elusive.

To explore the rules governing the propagation of single-gene perturbation effects, here we analyze the transcriptome profiles of single-gene deletion strains of *E. coli* and of CRISPRi-perturbed human and worm cells. Focusing first on *E. coli*, we show that more than a hundred genes change expression levels in most single-gene deletion strains. The analysis reveals sets of genes that maintain their expression level ratios (stoichiometry) against genetic perturbations, despite the lack of common gene regulatory and protein-protein interactions. By quantifying the strength of stoichiometry conservation between genes on a genome-scale, we uncover a simple rule that genes with high stoichiometry conservation with the deleted gene are more likely to be differentially expressed. Using large-scale Perturb-seq data, we further show that this relationship with stoichiometry conservation strength is also common in *Homo sapiens* and *Caenorhabditis elegans* cells, suggesting that this is a universal constraint conserved across organisms.

## Results

### Growth, transcriptomic, and Raman spectral changes caused by single-gene deletions

To investigate cellular state changes caused by single-gene deletions, we measured the growth curves, transcriptomes (RNA-seq), and Raman spectra of 21 single-gene deletion *E. coli* strains from the Keio collection and their parental strain *E. coli* K-12 BW25113 (*21*, *22*) (Fig. 1A, table S1). Growth curves were used as indicators of macroscopic phenotypic changes, while transcriptomes and Raman spectra were used as indicators of microscopic molecular changes such as expression states. The single-gene deletion strains were selected with reference to (*23*) to cover different growth phenotypes and to avoid bias of selected deleted genes towards specific gene functions (table S1).

**Fig. 1.**
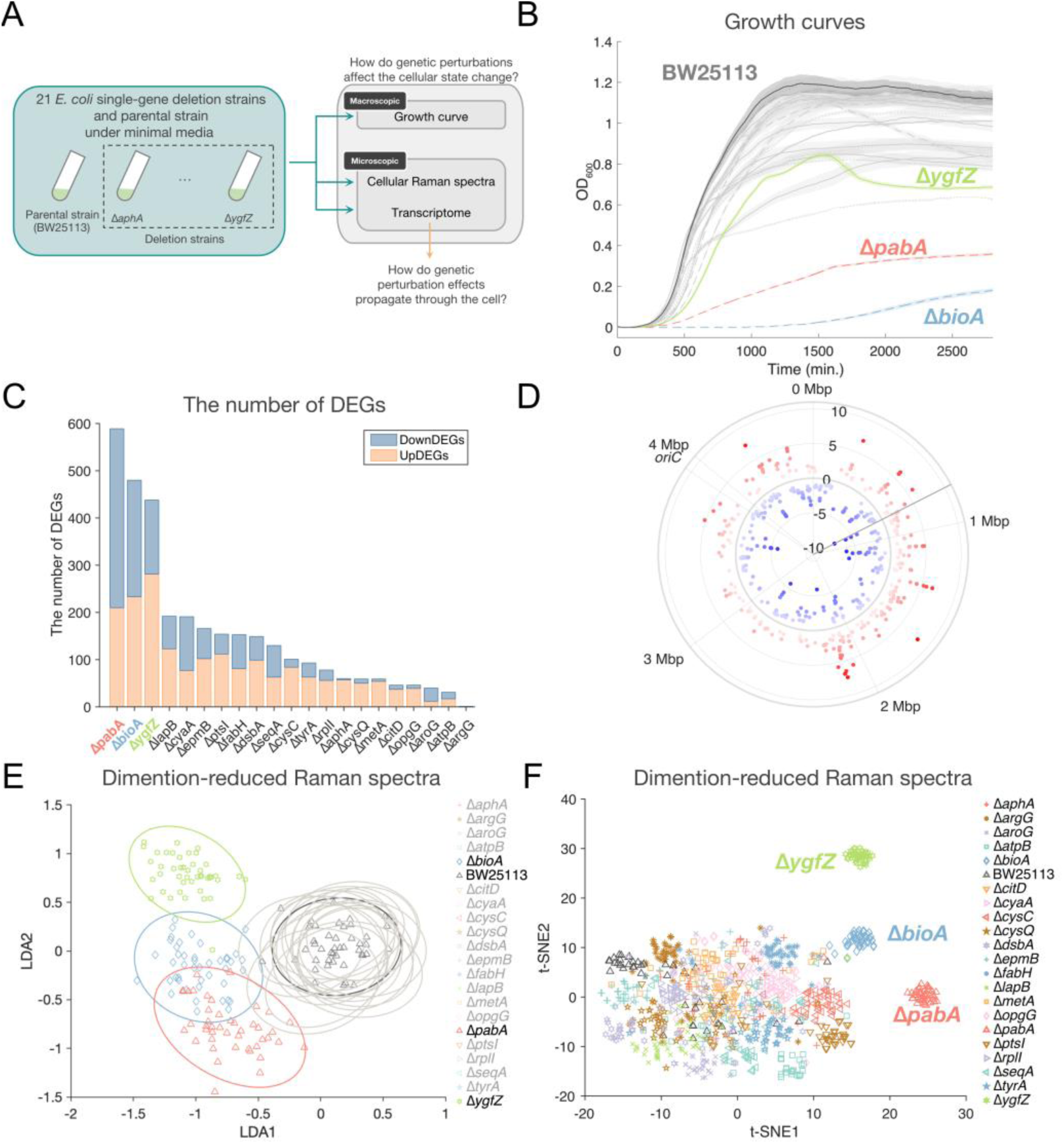
Changes in growth, transcriptome profile, and Raman spectrum caused by single-gene deletions. (**A**) Macroscopic and microscopic effects of single-gene deletions. (**B**) Growth curves of parental and single-gene deletion strains. The mean value for each replicate is shown as a thick line, with standard errors shown as shading above and below the mean value. The highlighted plots are the growth curves of parental strain and those of the three growth-defective strains. See Fig. S1 for the other deletion strains. (**C**) The number of DEGs of the deleted strains relative to the parental strain BW25113 (FDR<0.05, |log_2_FC|>0.7). UpDEGs are represented in red and downDEGs in blue. (**D**) Mapping of DEGs of the Δ*bioA* strain to *E. coli* chromosome. UpDEGs are colored in red, and downDEGs in blue, and radial coordinate values represent log_2_FC values. The position of the deleted gene is indicated by a solid grey line. (**E-F**) Cellular Raman spectra in LDA space (E) and t-SNE space (F). The low-dimensional (21 = #strain − 1) Raman spectra were calculated by PC-LDA and then embedded into two-dimensional space by t-SNE. Each point represents a spectrum from a single cell, and each ellipse shows the 95% concentration ellipse for each condition. The points for the strains other than the parental and the three growth-defective strains in the LDA space are shown in Fig. S17.

We first cultured all these 22 strains in minimal media supplemented with glucose and measured their growth curves (Fig. 1B, fig. S1). To quantitatively evaluate the differences in growth phenotype caused by gene deletions, we extracted four parameters from the growth curves: maximum growth rate, area under the curve (AUC), lag time, and maximum turbidity (maxOD) (table S2). The parental strain BW25113 had the highest growth rate and ranked the second highest AUC and maxOD, while most of the deletion strains had inferior growth parameters compared to the parental strain (Fig. 1B, fig. S1). Several strains displayed markedly different growth patterns. For example, the Δ*bioA* and Δ*pabA* strains had extremely low growth rates and maxOD; the Δ*ygfZ* strain had the third lowest growth rate and exhibited a characteristic decline in optical density after the transition from the exponential phase to the stationary phase; the Δ*cysC* strain, while not having a significantly reduced growth rate, had a notably low maxOD; and the Δ*dsbA* strain exhibited a continuous decline in turbidity after reaching maxOD (Fig. 1A, fig. S1). Given that some of these parameters were highly correlated (fig. S2A), we performed principal component analysis (PCA) to identify key attributes distinguishing strains with similar growth profiles (fig. S2B). The first two principal components explained approximately 95% of the variability in growth parameters among the selected strains (fig. S2C). Maximum growth rate, AUC, and maxOD were closely related to PC1, while lag time was most closely related to PC2 (fig. S2D).

To investigate the transcriptomic differences between the single-gene deletion strains, we next performed RNA-seq measurements on cells sampled from the exponential growth phase. We quantified the number of up- and down-differentially expressed genes (upDEG and downDEG) in each deletion strain compared to the BW25113 strain (Fig. 1C). The deletion strains with low growth rates (the Δ*bioA*, Δ*pabA* and Δ*ygfZ* strains) had a high number of DEGs (400–600), representing approximately 10–15% of the total *E. coli* genes. Most of the other deletion strains had around 100 DEGs, while some strains, such as the Δ*argG* strain, showed little to no change in gene expression. We mapped these DEGs onto *E. coli* chromosome, finding that most of them were not localized near the deleted genes but were distributed widely across the chromosomes, though some DEGs, such as those related to motility and chemotaxis, were clustered on chromosomes (Fig. 1D, and fig. S3). This implies that most single-gene deletions had genome-wide impacts on gene expression.

To determine whether the three strains with significant growth defects (the Δ*bioA*, Δ*pabA*, and Δ*ygfZ* strains) share common functional changes, we performed gene ontology (GO) enrichment analysis for all deletion strains and hierarchical clustering based on the similarity of enriched GO terms for both upDEGs and downDEGs (fig. S4). In the downDEGs of the strains with significant growth defects, we observed an enrichment of functions related to translation and amino acid transport, biosynthesis, and metabolic processes, indicating a down-expression of genes related to central functions required for growth. Since PabA physically interacts with proteins involved in amino acid biosynthesis, such as PheA, TrpABCDE, and TyrA (fig. S5), the functional enrichment of DEGs related to amino acid bioprocesses may reflect the effects of the lost protein-protein interactions between the products of these genes. However, BioA and YgfZ have no known regulatory and protein-protein interactions with genes involved in these central gene expression and metabolic processes. Moreover, PabA does not directly interact with genes involved in translation. Therefore, the down-expression of the genes central to growth in these strains could be mostly induced by some indirect and distant interactions in the intracellular networks.

GO enrichment analysis of the upDEGs indicated the induction of a cellular stress response in some of the deletion strains (fig. S4B). For instance, functions related to the colanic acid biosynthesis process were enriched in two of the strains with significant growth defects (the Δ*bioA* and Δ*ygfZ* strains) as well as in other strains (the Δ*fabH* and Δ*opgG* strains). Colanic acid is an exopolysaccharide produced by bacteria in the *Enterobacteriaceae* family, such as *E. coli*, and forms a capsule around a cell body (*24*). It is thought to contribute to biofilm formation and enhance bacterial survival in stressful environments, such as acidic conditions (*24*, *25*). The upDEGs of the Δ*pabA* strain were different from those of the Δ*bioA* and Δ*ygfZ* strains and enriched in the functions related to the glycolytic process and the nucleotide biosynthetic process (fig. S4B). Importantly, most of these upDEGs have no known regulatory and protein-protein interactions with the deleted genes. Therefore, it is likely that the up-expression of these genes is also induced by indirect and distant interactions.

Notably, a characteristic up-expression of motility-related genes was commonly observed in 9 of the 21 deletion strains (fig. S4B). Up-expression of motility-related genes in a significant fraction of the strains in the Keio collection has been reported previously and is likely caused by the rapid accumulation of secondary mutations that increase the expression of the major motility regulator (*26*). However, the up-expression of motility-related genes was absent in the three growth-defective strains (fig. S4B). Therefore, the extensive changes in the transcriptome profiles of these growth-defective strains cannot be explained by the up-expression of many motility-related genes.

To further investigate whether the observed large-scale changes in gene expression profiles probed by the DEG analysis result in changes in detectable global biomolecular composition, we measured the Raman spectra of the deletion strains at the single-cell level. Raman spectra are optical signatures that reflect the whole molecular compositions of cells (*27–29*). It has been shown that cellular Raman spectra are statistically correlated with transcriptomic and proteomic profiles and that these omics profiles can be inferred from the Raman spectra (*5*, *30–32*). Therefore, if the observed gene expression changes were indeed global and significant, the differences in cellular states of the deletion strains could be detected from their Raman spectra. We focused on the fingerprint region of the Raman spectra (700-1800 cm^-1^, fig. S6), where the signals from different biomolecules are concentrated (*27*, *33*). We reduced the dimensionality of the Raman spectra to 21 (=the number of strains – 1) dimensions using principal-component linear discriminant analysis (PC-LDA) and then t-distributed Stochastic Neighbor Embedding (t-SNE) were applied to dimension reduced Raman spectra (Figure 1E and 1F, fig. S7). The Raman spectra of the strains with significant growth defects (the Δ*bioA*, Δ*pabA*, and Δ*ygfZ* strains) formed distinct clusters separated from other deletion strains along the LDA1 axis in the low-dimensional LDA space, indicating the significant changes in their biomolecular profiles (Fig. 1E). The other deletion strains formed clusters overlapping with that of BW25113 in the LDA1-LDA2 space (fig. S7A, and S7B). However, all strains formed distinguishable clusters in t-SNE space (Fig. 1F). Thus, although the differences are smaller, the result suggests that the deletion strains other than the three growth-defective strains also possessed unique biomolecular profiles detectable by Raman spectra.

The results of the Raman spectral analysis are consistent with the observed growth and transcriptomic profiles, as significant correlations were found between the growth rate, the number of DEGs, and the LDA1 axis (fig. S8). Furthermore, the transcriptome profiles of the deletion strains were predictable from the Raman spectra (fig S9) (*5*, *30*). These results confirm from multiple perspectives that single-gene deletion has a global impact on their gene expression profiles and whole-cell molecular composition.

### DEGs are widespread in gene regulatory and metabolic networks, but the direction of expression changes is concerted in local units

To investigate the underlying principle for the propagation of gene deletion effects to other genes, we analyzed transcriptome data in combination with *E. coli* biological network information. We mapped the DEGs onto the *E. coli* gene regulatory network (GRN) obtained from RegulonDB v12.0 (*34*) and the metabolic network obtained from the KEGG database (*35*) to assess their relevance to the global gene expression changes (Fig. 2A, 2B, fig. S10, and S11). In the strains with significant growth-defects, many units of both regulons and transcription factors (TFs) showed differential expression changes (Fig. 2A, 2C, 2D and fig. S10), confirming global effects of the single-gene deletion. We examined whether the DEGs were restricted to the genes that shared TFs with the deleted gene, but found that a significant proportion did not share the same TFs (Fig. 2E). Furthermore, we calculated and compared the network distance between the DEGs and the deleted genes, i.e., the shortest path length between these genes (fig. S12-S16) and found that both upDEGs and downDEGs are not restricted to the vicinity of the deleted genes. However, this result does not imply that the DEGs are independent of GRNs. In fact, the direction of expression changes (up- or down-expression) within each regulon or pathway was often concordant (Fig. 2A, 2B, fig. S10, and S11); DEGs in operons and pathways often showed expression changes in the same direction at rates significantly higher than those predicted by random assignment of upDEGs and downDEGs (operon: *p* < 0.001 for all strains, pathway: *p* < 0.05 for all strains expect for the Δ*atpB* strain (*p* = 0.073)); Fig 2A and fig. S17). Thus, the changes in gene expression levels of DEGs tended to be concerted within local units of the GRN.

**Fig. 2.**
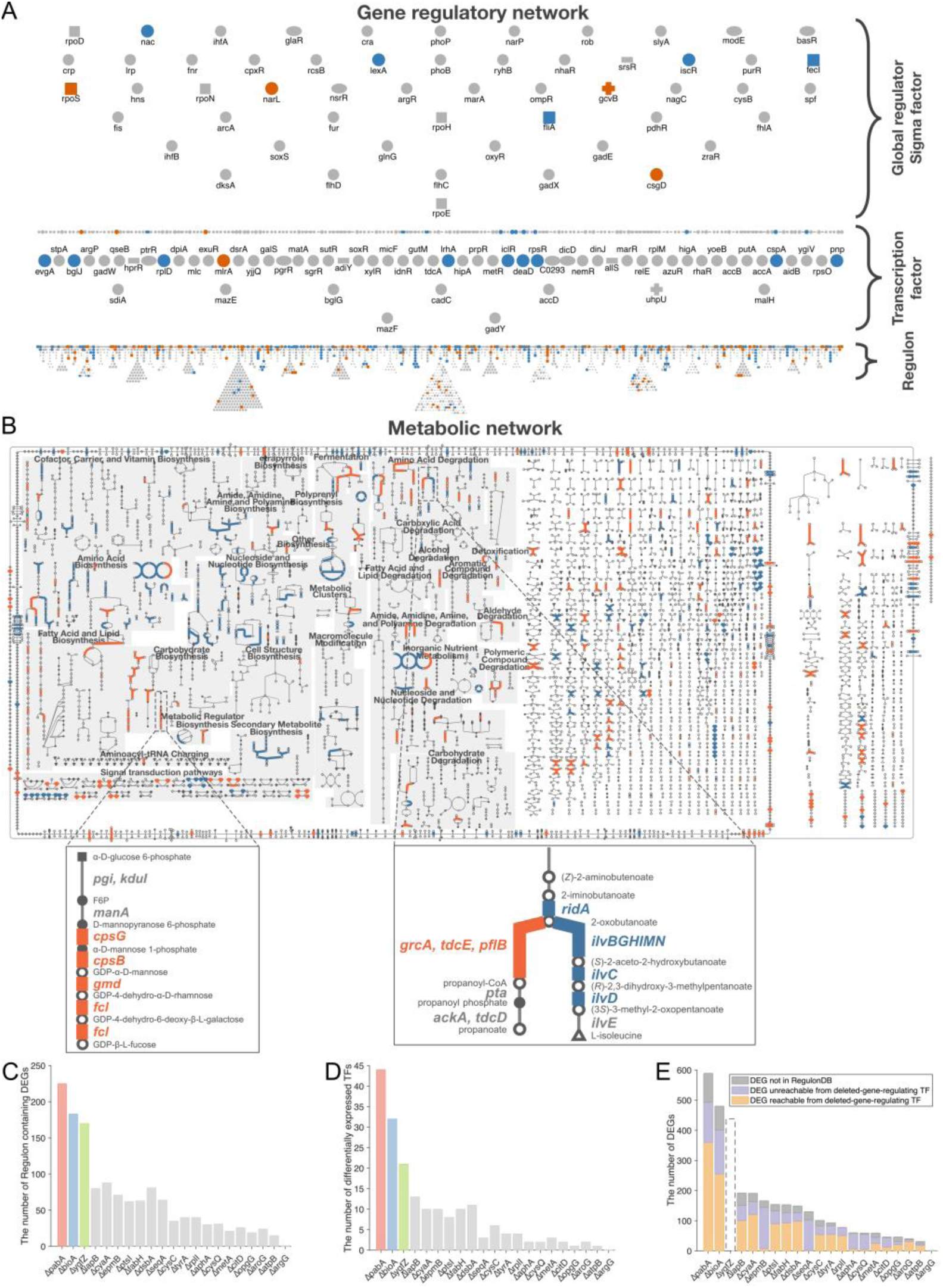
Global propagation of single-gene deletion effects. (**A**) Mapping the DEGs of the Δ*bioA* strain to the *E. coli* gene regulatory network. UpDEGs are colored in red, and downDEGs in blue. Mapping was performed using the Regulatory Overview from EcoCyc (*36*). (**B**) Mapping of DEGs of the Δ*bioA* strain to the *E. coli* metabolic network. Pathways involving upDEGs are colored in red, and downDEGs in blue. Mapping was performed using the Regulatory Overview from EcoCyc (*36*). The magnified view shows the GDP-mannose biosynthesis, GDP-L-fucose biosynthesis I pathway (left), in which all DEGs are up-expressed, and the branching L-threonine degradation I, L-isoleucine biosynthesis I pathway (right), in which the DEGs show concerted changes in each branch but the directions of the changes are opposite between the branches. (**C**) Histogram of the number of regulons containing DEGs. X axis is sorted by the number of DEGs. (**D**) The number of differentially expressed TFs in the deletion strains. TFs are defined as genes that have at least one regulating target gene in the gene regulatory network of EcoCyc. (**E**) The proportion of DEGs that share the same TFs with the deleted gene. DEGs regulated by TFs that regulate the deleted gene are shown in orange; DEGs not regulated by TFs that regulate the deleted gene are shown in purple; and DEGs not included in the database used (RegulonDB (*34*)) are shown in gray. *ygfZ* is not included in RegulonDB and is shown as an empty column.

Mapping DEGs onto the metabolic network again characterizes the widespread distribution of DEGs in different pathways (Fig. 2B). Therefore, the affected genes are not limited to neighboring pathways in which the deleted gene was involved. Again, this result does not imply that the DEGs are independent of the metabolic network. In fact, although we can find DEGs with the expression changes in different directions in branched metabolic pathways, the directions of expression changes were almost always consistent in unbranched metabolic pathways (Fig 2B, and fig. S11). Therefore, the changes in gene expression levels of DEGs involving the same local metabolic pathways are concerted.

### Stoichiometry conservation of functionally related genes against gene deletion perturbations beyond local operons

The observed expression changes of DEGs in the same directions in local regulons or metabolic pathways (Fig. 2A, 2B, fig. S10, and S11) may hint at a principle underlying the observed global gene expression changes in the deletion strains. To further investigate these locally coordinated expression changes in metabolic pathways, we next examined not only the directions but also the magnitudes of gene expression changes in local units of these intracellular networks.

As an example, we first examined the expression level of the genes in the L-tryptophan biosynthesis pathway (*trpABCDE*), all of which are in the *trp* operon (Fig. 3A, and 3B). The expression levels of these five genes varied widely among the deletion strains, but the directions and magnitudes of the expression changes were highly concerted in all strains (Fig. 3A). Comparison of the normalized expression levels of these five genes shows almost identical patterns of change in all strains except the Δ*pabA* strain, which had a significantly lower expression level of the *trp* genes (Fig. 3B). Therefore, the ratio of expression levels (stoichiometry) of these five genes is strongly conserved across different gene deletions.

**Fig. 3.**
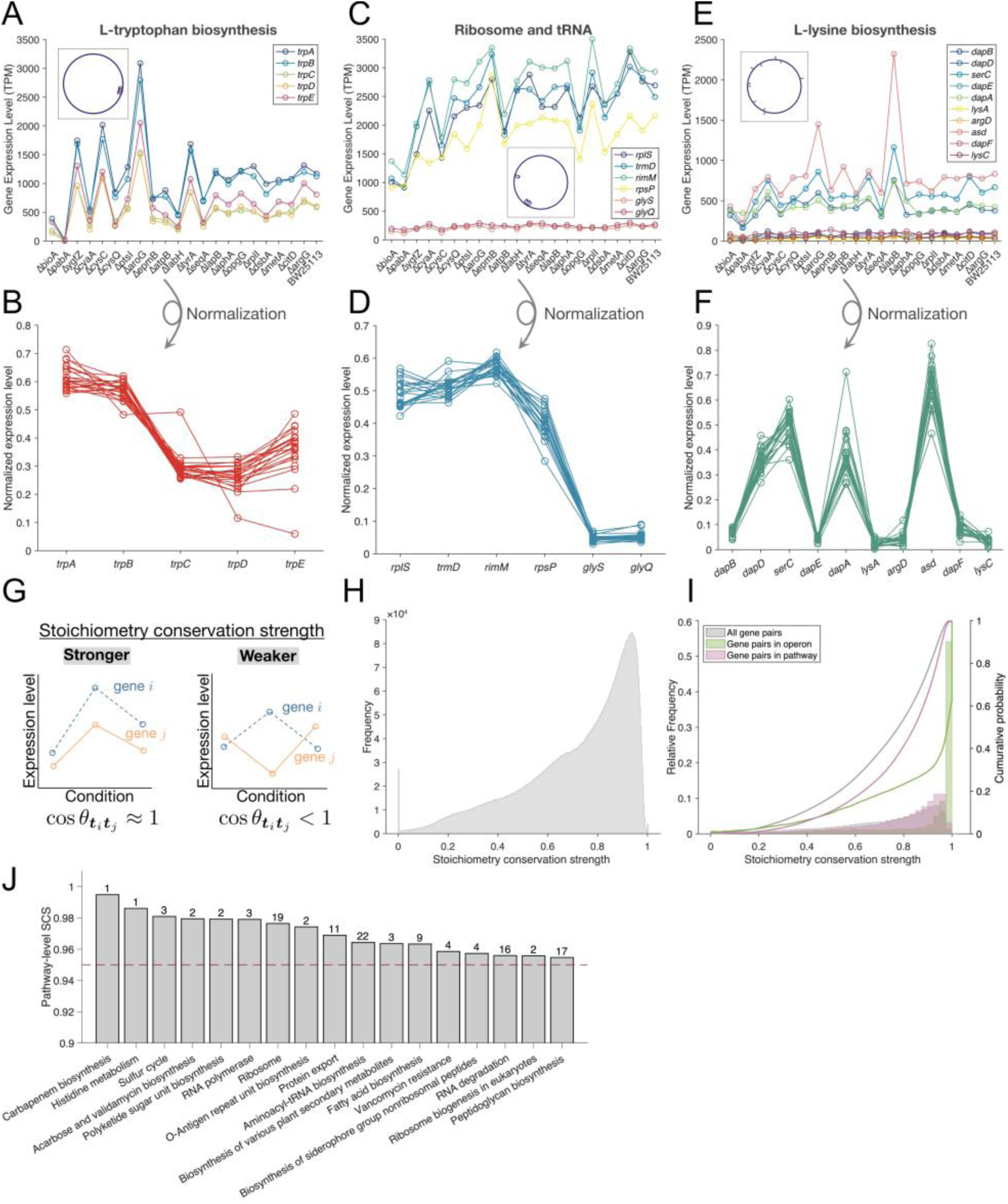
Stoichiometry conservation against single-gene deletion perturbations within operons and pathways. (**A**) Gene expression levels (TPM) of five genes involved in the L-tryptophan biosynthesis pathway (*trp* opron). The inset is chromosomal location of these five genes. The x-axis is sorted by ascending growth rates. (**B**) Expression levels of five genes involved in L-tryptophan biosynthesis pathway (*trp* opron) under 22 conditions normalized by L2 norm. (**C**) Gene expression levels (TPM) of six genes involved in the *rpsP-rimM-trmD-rplS* and *glyQS* operons. These two operons are 0.98 Mbp apart on the chromosome and share no common transcription factors. The inset is chromosomal location of these six genes. The x-axis is sorted by ascending growth rates. (**D**) Expression levels of six genes involved in *rpsP-rimM-trmD-rplS* and *glyQS* operon under 22 conditions normalized by L2 norm. (**E**) Gene expression levels (TPM) of ten genes involved in the L-lysine biosynthesis pathway. These ten genes share no common transcription factors. The inset is chromosomal location of these ten genes. The x-axis is sorted by ascending growth rates. (**F**) Expression levels of ten genes involved in the L-lysine biosynthesis pathway under 22 conditions normalized by L2 norm. (**G**) Schematic of stoichiometry conservation relations. For each gene species, its expression vector is considered, whose elements represent the gene expression levels in different strains. The cosine similarity between the expression vectors of two gene species is close to 1 if the expression stoichiometry between genes is strongly conserved across conditions but is less than 1 if the expression patterns are inconsistent. (**H**) Histogram of stoichiometry conservation strength of all gene pairs calculated based on the expression level changes in the 22 *E. coli* strains. (**I**) Histograms of stoichiometry conservation strength of gene pairs restricted to those in operon (green) and metabolic pathway (pink) compared with the distribution of all gene pairs. The left y-axis is for the relative frequency distributions (histograms), and the right y-axis is for the cumulative distributions (lines). (**J**) Pathways with high pathway-wise mean SCS up to the pathway above the threshold of 0.95 (red line). The number above the bar indicates the number of operons involved in each pathway.

The stoichiometry conservation of the *trp* genes may be trivial because these five genes are under the control of the same promoter and are transcribed together. However, we also found several sets of functionally related genes whose direction of expression level change and mutual stoichiometry were consistent across multiple operons (e.g., ribosome and tRNA genes, Fig. 3C, and 3D). In addition, such strong conservation of mutual stoichiometry was also found for a set of genes that do not share operons, such as those involved in L-lysine biosynthesis pathways (Fig. 3E, and 3F). Thus, stoichiometry conservation is not limited to units with shared transcriptional control, but is a property that exists beyond the local gene regulatory architecture.

To evaluate the prevalence of stoichiometry conservation among genes in operons and local metabolic pathways systematically, we quantified the stoichiometry conservation strength (SCS) of a gene pair against gene deletion perturbations by cosine similarity of their expression vectors (Fig. 3G) (*5*), i.e.,

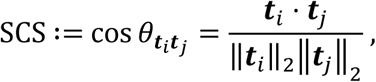

where 𝒕_𝑖_: = (𝑡_𝑖1_, 𝑡_𝑖2_, …, 𝑡_𝑖𝑛_)^⊤^ is the expression vector of gene 𝑖, whose elements represent its expression levels in the different deletion strains (𝑛 = 22); and || ⋅ ||_2_ represents the 𝐿^2^norms. The SCS takes values between 0 and 1 and will be closer to 1 when two genes strongly maintain their stoichiometry across different deletion strains.

The distribution of SCS for all gene pairs was highly skewed toward high values (Fig. 3H), as previously observed for environmentally perturbed *E. coli* and cells (*5*). Thus, there is a general genome-wide trend for genes to maintain mutual stoichiometry against both genetic and environmental perturbations. We also calculated the SCS distributions by restricting the gene pairs to those within the same operons and to those within the same pathways, and found that these distributions are even more skewed toward higher SCS values (Fig. 3I, *p* < 10^-15^ for both operon and pathway by the Brunner-Munzel test). Thus, the genes in the same operon or local pathways tend to maintain their stoichiometry more strongly.

We also calculated the mean SCS for each metabolic pathway registered in the KEGG database (*35*, *37*) (Fig. 3J). Interestingly, some of the pathways with high mean SCS, such as those related to ribosome, protein export, and aminoacyl-tRNA biosynthesis, were composed of many operons (Fig. 3J). Therefore, the observed stoichiometry conservation is likely the product of multi-level regulation and the coordination of expression levels beyond operons (*5*).

### DownDEGs in the growth-defective deletion strains are high SCS genes with the deleted genes

The prevalence of stoichiometry-conserving gene pairs beyond local gene regulatory units led us to ask whether genes that strongly maintain stoichiometry with a deleted gene are more likely to be differentially expressed (Fig. 4A). To test this, we compared the SCS with the deleted genes between all genes and DEGs. Here, we focused on the three growth-defective strains (the Δ*bioA*, Δ*pabA*, and Δ*ygfZ* strains) that had many DEGs and calculated the SCS with these deleted genes using the transcriptome data of the other 19 strains.

**Fig. 4.**
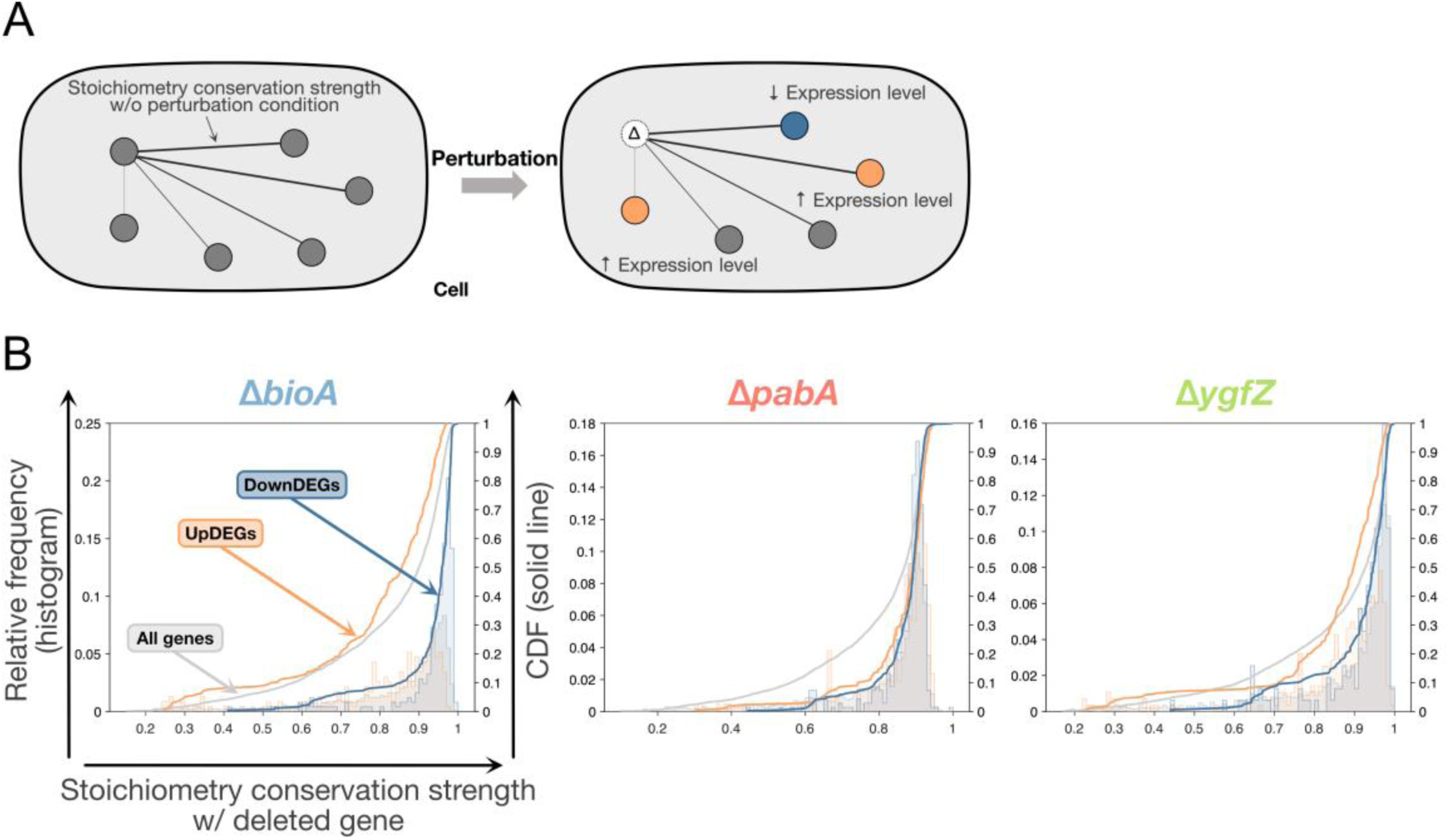
Enrichment of downDEGs in the gene fraction with strong stoichiometry conservation with the deleted genes. (**A**) Schematic of the analysis. Here, we examined the relationship between the genome-wide gene SCS with the deleted gene and the resulting gene expression changes due to the deletion. (**B**) Propagation of gene deletion effects to genes that conserve high expression stoichiometry with the deleted gene. The x-axis represents SCS; the closer to 1, the higher the stoichiometry conservation strength with the deleted genes. The left y-axis is for the histograms showing the relative frequency of the SCS of upDEGs, downDEGs and all genes (orange, blue and grey, respectively), and the right y-axis is for the solid lines that represent cumulative distribution functions of the SCS distributions.

The result showed that downDEGs had significantly higher stoichiometry conservation with the deleted genes compared to other genes across all three strains (Fig. 4B, 𝑝 < 10^−15^ for Δ*bioA* and Δ*pabA*; and 𝑝 < 10^−4^ for Δ*ygfZ*, by Brunner-Munzel test (*38*)). On the other hand, upDEGs showed no consistent pattern across the strains with respect to the strength of stoichiometry conservation: the stoichiometry conservation of the upDEGs was high in the Δ*pabA* strain, but not in the Δ*bioA* and Δ*ygfZ* strains. These results show that at least the downDEGs are the genes that had strong stoichiometry-conserving relationships with the deleted genes.

### Generality of propagation of genetic perturbation effects through stoichiometry conservation architecture

To further extend the analysis and investigate the general relevance of stoichiometry conservation relations to global gene expression changes caused by genetic perturbations, we analyzed two gene expression profile datasets of genome-wide genetic perturbations: Genome-Wide Perturb-Seq (GWPS) data from human chronic myeloid leukemia (K562) cells (*13*) and Worm Perturb-Seq (WPS) data from *C. elegans* cells (*39*, *40*). Perturb-seq is a CRISPR-based, high-throughput screening method that uses single-cell RNA-sequencing to assess comprehensive gene expression profiles following genetic perturbation (*15*, *41*).

Although CRISPRi differs from gene deletion, both schemes of genetic perturbation result in reduced gene expression. In both the GWPS and WPS datasets, we focused on the data from perturbation conditions that resulted in large transcriptional changes (perturbation with ≥ 200 DEGs for GWPS data and ≥50 DEGs for WPS data, respectively) and the unperturbed control. To evaluate the effect of inactivating a particular gene, we first excluded the transcriptome data of the sample with its particular gene inactivated and calculated the SCS between the perturbed gene and other genes using the remaining data. We then examined whether the DEGs in this perturbation condition had higher SCS with the perturbed gene.

The result showed that, in most genetic perturbation conditions, both upDEGs and downDEGs had high SCS with the perturbed gene in both human and worm cells (Fig. 5A and fig. S18). The enrichment of genes with high SCS genes in upDEGs and downDEGs was observed across a wide range of genetic perturbation conditions with varying numbers of DEGs (Fig. 5A and fig. S18).

**Fig. 5.**
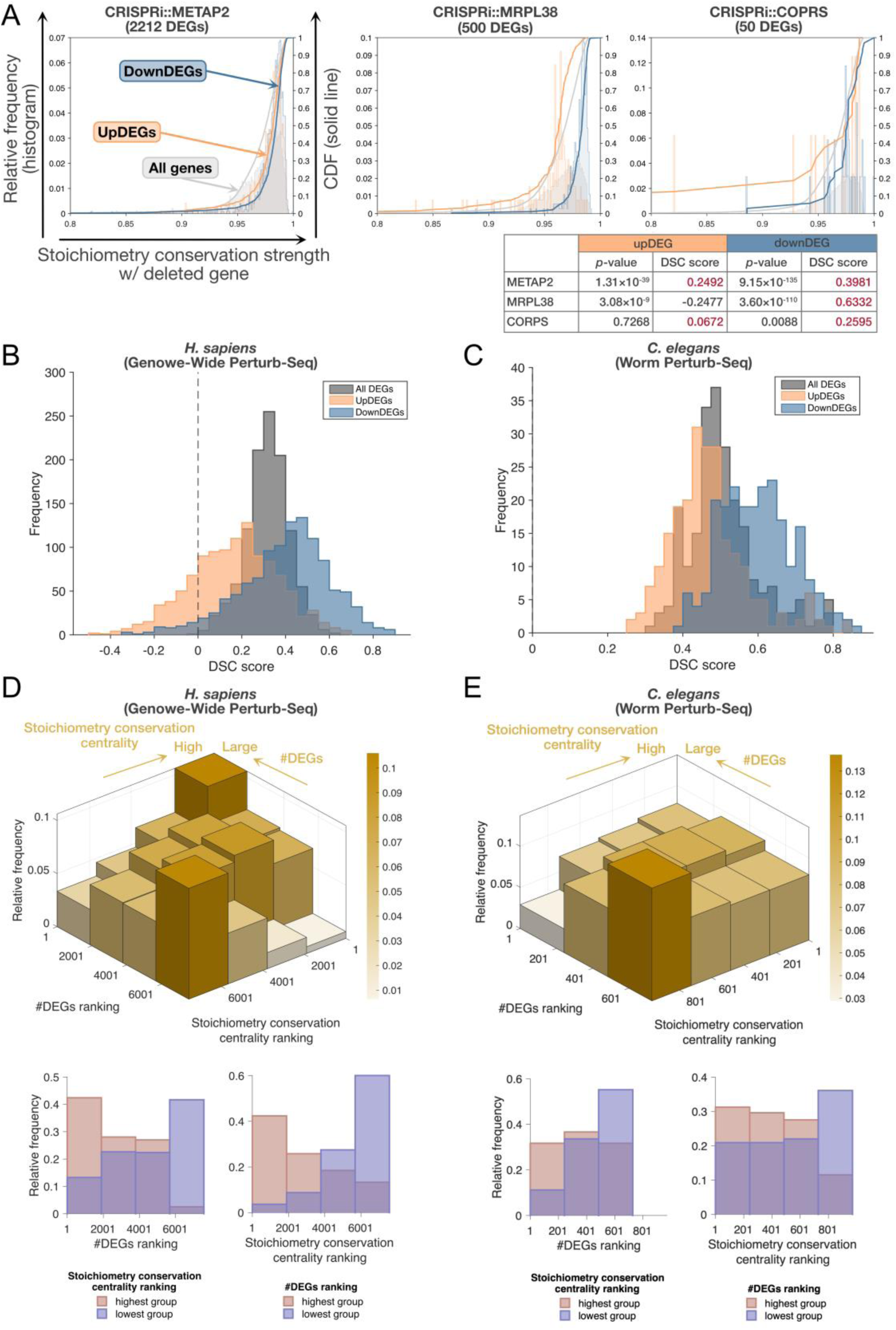
Generality of propagation of genetic perturbation effects through stoichiometry conservation relationship. (**A**) Propagation of genetic perturbation effects to genes that conserve stoichiometry with the perturbed gene in human. The x-axis represents SCS between the perturbed gene; the closer to 1, the higher the stoichiometry conservation relation strength of the deleted genes. The left y-axis is for the histograms showing the relative frequency of the stoichiometry conservation relation strength of upDEGs, downDEGs and all genes, and the right y-axis is for the solid lines that represent cumulative distribution functions of the SCS distributions. The table summarizes the results of *p*-value and DSC score, which were evaluated using Brunner-Munzel test to evaluate whether the SCS distribution of upDEGs and downDEGs is higher than the SCS distribution of all genes in each perturbation. Positive DSC scores are highlighted in red. (**B, C**) Histogram of DSC scores in human (B) and in *C. elegans* (C). The DSC score is greater than 0 when DEGs tend to conserve stoichiometry with with the perturbed gene, while it is less than 0 when DEGs are less likely to conserve stoichiometry with the perturbed gene. (**D-E**) Correlation between stoichiometry conservation centrality and the number of DEGs in the human cells (D) and in the *C. elegans* cells (E). The plots in the lower left of the 3D histograms compare the distributions of the DEG number rankings between the highest and the lowest groups of the stoichiometry conservation centrality rankings; the plots in the lower right compare the distributions of stoichiometry conservation centrality rankings between the highest and the lowest groups of DEG number rankings. The gene fraction with many DEGs tends to have higher stoichiometry conservation centrality (*H. sapiens*: 𝑝 < 10^−15^; *C. elegans*: 𝑝 < 10^−12^, the one-sided Brunner-Munzel test). The gene fraction with high stoichiometry conservation centrality tends to have more DEGs (*H. sapiens*: 𝑝 < 10^−15^; *C. elegans*: 𝑝 < 10^−9^, the one-sided Brunner-Munzel test).

To investigate whether this trend is consistent across perturbations, we introduced a metric called the DEG stoichiometry conservation (DSC) score. The DSC score quantifies the overall degree of the distribution shift of SCS between the DEGs and the perturbed gene based on the relative treatment effect in the Brunner-Munzel test (see Materials and Methods). A positive DSC score means that the DEGs have stronger stoichiometry conservation relations with the perturbed gene compared to all genes, whereas a negative DSC score means the opposite.

We found that both upDEGs and downDEGs exhibited significantly higher DSC scores in most perturbation conditions in both organisms (Fig. 5B, and 5C). The downDEGs had overall higher DSC scores compared to the upDEGs. In the GWPS datasets, we found a significant fraction (20.6%) of the perturbation conditions where upDEGs showed negative DSC scores (Fig. 5B), as observed for the upDEGs in *E. coli* (Fig. 4B).

Because it evaluates the shift of the entire SCS distribution, the DSC score tends to underestimate the trends in the left tail, i.e., the characteristics of the genes with low SCS with the deleted gene. Therefore, we examined the expression changes of the low SCS genes and found that they are more likely to be up-expressed when the corresponding gene is inactivated (fig. S19). This result suggests that genes that have low SCS with the perturbed gene (i.e., genes whose expression levels can change without the constraint of maintaining stoichiometry with the perturbed gene) are more likely to be up-expressed when they change the expression level. Since downDEGs are enriched in the higher DSC score fractions than upDEGs, the enrichment of up-expressed DEGs in the low SCS fraction could be the consequence of compensating for the expression changes in the high SCS genes.

If genetic perturbation effects propagate to other genes through stoichiometry-conserving relationships between genes, it is plausible that genes that maintain stoichiometry with more genes will have more DEGs when their expression is inactivated. To test this speculation quantitatively, we calculated the stoichiometry conservation centrality of each gene by summing its SCS with all other genes and found that centrality is indeed positively correlated with the number of DEGs when its expression is perturbed by CRISPRi in both the GWPS and WPS datasets (Fig. 5D and 5E).

Taken together, these findings suggest that the effects of genetic perturbations are propagated to genome-wide genes through a stoichiometry-conserving architecture, and that this feature is universal across different organisms and types of genetic perturbations.

## Discussion

In this study, we investigated how genetic perturbation effects propagate within cells. Our results highlight the stoichiometry conservation relations between the perturbed gene and others. Specifically, when a gene is perturbed, genes that maintain strong stoichiometry relations with the perturbed gene tend to be differentially expressed. This feature is found not only in *E. coli* but also in human and *C. elegans* cells. Therefore, stoichiometry conservation relation constrains how genome-wide gene expression state changes in response to genetic perturbations.

Our analysis of *E. coli* indicated that the stoichiometry conservation between genes correlates with the architecture of gene regulatory networks, such as operons and transcription units. However, stoichiometry conservation was observed beyond such gene regulatory units in *E. coli*. Furthermore, similar stoichiometry conservation architecture existed even in human cells, where most genes are expressed monocistronically. Therefore, the stoichiometry conservation of genome-wide genes is likely the product of more complex and multi-level constraints in cells beyond transcriptional control.

Stoichiometry conservation relation likely plays a crucial role in maintaining cellular function and physiology (*5*, *42–44*). For example, in-pathway and functionally analogous enzyme stoichiometry is quantitatively conserved throughout evolution, not only within bacteria but even in eukaryotes (*42*). Quantitative analysis combining proteome and Raman spectra data from *E. coli* revealed a low-dimensional hierarchical stoichiometry-conserving structure in the proteome, underpinning principles that facilitate cellular homeostasis and adaptation. Furthermore, antibiotics targeting ribosomes disrupt ribosomal protein balance and halt growth in *E. coli*, but the balance is restored along with growth recovery during physiological adaptation (*45*). Theoretical analysis shows that exponential cell growth requires stoichiometry conservation within pathways (*44*). These findings suggest that stoichiometry conservation contributes to optimal cellular function, but our study suggests that stoichiometric conservation may also be constraint on the expression profile changes caused by genetic perturbation.

An intriguing question is whether the stoichiometry conservation architecture identified in our study holds true for other types of perturbations, such as overexpression. Overexpression plays a significant role in stem cell induction, differentiation, and microbial drug resistance. Genome-wide profiling of genes that enhance stress resistance in *Saccharomyces cerevisiae* under stress condition through adaptive overexpression (*46*) and systematic analyses of transcription factor overexpression along the EMT (epithelial-mesenchymal transition) axis have been conducted (*47*). However, genome-scale expression profiles for overexpression remain limited compared to inactivation studies, which currently poses a challenge for the systems-level analysis of overexpression effects.

Many organisms are known to possess the mechanism to compensate for expression changes caused by copy number changes of genes, e.g. due to aneuploidies of genomes (*48*). The observed global gene expression changes due to genetic inactivation and the ubiquity of stoichiometry conservation between genes may be related to the mechanisms responsible for this dosage compensation.

From a technical point of view, an advantage of the metric, stoichiometry conservation strength (SCS), introduced here is that it can be evaluated only from gene expression data without relying on the underlying biological networks that caused such coordination of gene expression levels. In general, our knowledge of biological networks such as gene regulatory networks and protein-protein interaction networks is still incomplete. Moreover, the resulting genome-wide gene expression profiles under certain conditions may well be the consequences of multi-level regulation. Therefore, the analysis of SCS, which does not explicitly require information about the underlying regulatory networks that caused the stoichiometry conservation, can be easily applied to different organisms, which could facilitate the characterization of universal constraints that exist beyond simple model organisms.

Despite the relevance of genome-wide stoichiometry conservation architecture to the gene expression changes by genetic perturbation, predicting resulting gene expression profiles in response to specific genetic perturbations remains a challenging task. Recent generative pretraining models, such as scGPT (*49*), has proven its effectiveness, but emerging evidence suggests that simple linear or baseline models can perform as well as or even better than scGPT (*50*, *51*). Future studies will clarify whether the stoichiometry conservation architecture can help improve predictions of genetic perturbation effects and provide systems-level understanding on how genetic perturbations give rise to specific phenotypic consequences.

## Acknowledgments

We thank Christine Jacobs-Wagner and Manuel Campos for sharing growth and physiology data of the *E. coli* strains in the Keio collection; the members of the Wakamoto Lab for discussion. This work was supported by JST CREST Grant Number JPMJCR1927 (Y.W.) and JPMJCR18S3 (K.O.); JST ERATO Grant Number JPMJER1902 (Y.W.); Japan Society for the Promotion of Science KAKENHI Grant Number 24H00552 (Y.W.), 19J22448 (K.F.K.), 24K02068 (K.O.), 24H01393 (A.H.O.) and 23K04984 (A.H.O); and AMED Grant JP22gm1610007 (K.O.).

## Materials and Methods

### *E. coli* Strains and Culture Medium

21 *E. coli* single-gene deletion mutants from the Keio collection (*21*, *22*) and their parental strain BW25113 were used. The 22 strains are listed in table S1. Cells were grown in M9 minimal medium supplemented with 0.2% glucose and 0.1% casamino acids at 30°C, which is the same culture medium in (*23*). The chemical composition of our M9 medium was also the same as that in (*23*).

### Cultivation

Cells were taken from a glycerol stock and cultivated for about 15 hours. The cultivated cells were then diluted into pre-warmed culture medium at an OD_600_ of around 0.01 and grown up to exponential phase prior to growth curve measurements, RNA extraction, and cellular Raman spectra measurements. Δ*bioA* reached exponential phase at an OD_600_ of round 0.1, Δ*pabA* around 0.2, and the other 20 strains around 0.3.

Cultivation was conducted with 2 mL medium in a glass test tube with a diameter of 16.5 mm and a length of 165 mm under reciprocal shaking at 200 r/min.

### Growth Curve Measurements and Analysis

Cells in exponential phase were diluted into pre-warmed culture medium at an OD_600_ of around 0.001 and cultivated. Growth curves were obtained by continuously recording turbidity every five minutes (ODBox-C, TAITEC CORPORATION). Three to eight biological replicates were obtained per strain.

Cultivation for growth curve measurements were conducted with 5 mL culture media, not 2 mL, due to a requirement of the turbidity recording device.

Four growth parameters (maximum growth rate, maximum OD, area under curve, and lag time) were extracted from the growth curves. Maximum growth rate was determined as the average of three replicates of maximum growth rate estimated using established methods. (*52*). maximum OD, area under curve, and lag time were defined as maximum OD value, the area under the growth curve from the start of measurement to 2800 minutes, the time (min.) until the OD value first exceeded 0.007, respectively.

### Raman Measurements and Preprocessing of Spectra

Cells in exponential phase were washed three times with 0.9% aqueous solution of NaCl, and 5 µL of the suspension was placed on a synthetic quartz slide glass (TOSHIN RIKO) and dried. Raman spectra of cells were measured with a Raman microscope, where a custom-built Raman system (STR-Raman, AIRIX) was integrated into a microscope (Ti-E, Nikon). Excitation light was generated by a 532nm continuous-wave diode-pumped solid-state laser (Gem 532, Laser Quantum). Light from the laser oscillator was transmitted by mirrors in this research. A 100× and NA = 0.9 air objective lens (MPLN100X, Olympus) was used. Raman scattered light was collected by an optical fiber, transmitted to a spectrometer (Acton SP2300i, Princeton Instruments) and dispersed by a 300 gr/mm grating. Dispersed light was projected onto an image sensor of an sCMOS camera (OrcaFlash 4.0 v2, Hamamatsu Photonics). The sCMOS camera was water-cooled at 15°C to reduce dark noise. The optical setup is the same as in (*5*). The exposure time for each single cell was 10 s. Randomly selected 15 to 16 cells were measured per strain per replicate. Raman spectrum of background was measured for each cell with 10 s exposure in an area close to a targeted cell where neither cells nor NaCl crystals existed. The laser power at the sample stage was 26 mW. The measurement system and processes were controlled by using μManager 1.4 (*53*) and a plugin we made. Three biological replicates were obtained per strain.

Obtained spectral images were preprocessed before the following analysis. See (*5*) for detail.

### Principal Component-Linear Discriminant Analysis (PC-LDA) and t-SNE of Raman Spectra

To reduce dimensions of cellular Raman spectra, we conducted principal component-linear discriminant analysis (PC-LDA). PC-LDA is a supervised classification technique that combines principal component analysis (PCA) and linear discriminant analysis (LDA) to find the most discriminatory bases while avoiding over-fitting. PCA was first applied to the original Raman spectra and principal components were selected that explained 98% of the variance in the original Raman spectra. Then, LDA was applied to selected principal components to extract the 21 (#strain – 1) most discriminatory bases by maximizing the ratio of the between-strain variance to the sum of within-strain variances in the lower dimensional space.

### Raman-Omics Correspondence by Partial Least Squares Regression (PLS-R)

To examine whether different transcriptomes could be estimated by Raman spectra in the LDA space, we conducted partial least squares regression (PLS-R). See (*30*) for detail.

To evaluate the correspondence between dimension reduced Raman spectra and transcriptome data, we conducted a leave-one-out cross-validation (LOOCV). Statistical significance of the correspondence was tested with permutation tests (*54*). In permutation test, PLS-R was run on the correct Raman spectral and transcriptome data pairs to learn the correspondences. Group labels were randomly shuffled to generate 10,000 false data sets. The empirical *p*-value was computed as <u>(𝑏 +</u> *<u>1</u>*<u>)/(𝑚 +</u> *<u>1</u>*) where 𝑏 is the number of permutations that gave the values equal to or lower than the original prediction error value, and 𝑚 is the total number of permutations (10,000). See (*30*) for detail.

### RNA Sample Preparation and RNA Sequencing

Cells in exponential phase were used for RNA extraction. First, RNAprotect Bacteria Reagent (QIAGEN) was added to cell cultures in exponential phase, and then the total RNA was extracted using RNeasy Mini Kit (QIAGEN). Further DNA removal was conducted using Recombinant DNase I (Takara Bio). Sequence libraries were prepared using KAPA RNA HyperPrep Kit (Kapa Biosystems). NEBNext Multiplex Oligos for Illumina (New England Biolabs) was used instead of KAPA adapters (Kapa Biosystems). 150 bp paired-end sequencing was conducted with HiSeq X (Illumina) by Macrogen Japan Corp. Two biological replicates were obtained per strain.

At the time of these experiments, reagents to remove bacterial rRNA were temporarily not commercially available at all. Therefore, we quantified total RNA using sufficiently large number of reads.

### RNA-seq Data Processing

The obtained sequences were trimmed, mapped to the E. coli BW25113 genome sequence (accession no. NZ_CP009273) with STAR (v) (*55*) using following parameters; –runMode alignRead –outSAMtype BAM SortedByCoordinate –rumThreadN 4 – quantMode TranscriptomeSAM. The BAM files output by STAR were read-counted by RSEM (v), counting the reads and calculating the TPM values. Differential expression analysis relative to parental strain were conducted with EdgeR (v.4.0.16) (*56*) using the following threshold; FDR<0.05, |logFC|>0.7.

### DEG Enrichment Analysis

Differentially expressed genes (DEGs) identified from RNA-seq analysis were subjected to GO (GOTERM_BP_DIRECT) enrichment analysis using the Database for Annotation, Visualization, and Integrated Discovery (DAVID) (https://david.ncifcrf.gov) (*57*, *58*). The DEGs were uploaded as a gene list. Enrichment significance was assessed using a modified Fisher’s exact test (EASE score), and *p*-values were corrected for multiple testing using the Benjamini-Hochberg method. Functional categories with an adjusted *p*-value of <0.05 were considered significantly enriched.

### Network Distance and Centrality Analysis

Network distances from each perturbed gene to DEGs were calculated using the MATLAB “distances” function. These network distances and centralities were calculated for publicly available gene regulatory networks (RegulonDB v12.0 (*34*)), protein-protein interaction networks (STRING-db (*59*)), and functional networks (EcoliNet (*60*)).

### Stoichiometry Conservation Relation Analysis

We extracted groups of genes with high stoichiometry conservation strength from gene expression profile in a systematic way. Consider a vertical vector that represents the expression level (TPM) of gene 𝑖 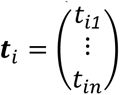. Each component of this vector represents the expression level of expression in each strain of gene 𝑖. This time, 𝑛 = *22* because 22 strains were used for the parent and deletion strains. Using this, we first evaluated the similarity of the gene expression pattern of all gene combinations with cosine similarity. The cosine similarity between genes 𝑖 and 𝑖𝑗 can be calculated as

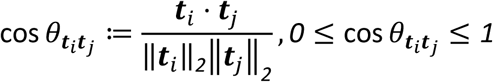

which is called stoichiometry conservation strength. cos 𝜃_𝒕𝑖𝒕𝑗_ takes a maximum value of 1 if and only if 𝒕_𝑖_ and 𝒕_𝑗_ are oriented in the same direction in condition space, i.e. the expression stoichiometry between gene 𝑖 and gene 𝑗 does not change when the conditions are changed. In other words, this can be regarded as a quantitative indicator of how much the expression stoichiometry is conserved when the condition changes.

We evaluated stoichiometry conservation strength between genes in each operon/pathway by operon-/pathway-level SCS (stoichiometry conservation strength) defined as below

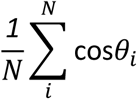

where 𝑁 is the number of gene pairs in each operon/pathway, and cos𝜃_𝑖_ is stoichiometry conservation strength of gene pair 𝑖. For calculation operon-level SCS, we excluded monocistronic operons, and operons with at least one zero expression gene. For calculation pathway-level SCS, we excluded genes not included in transcriptome data, and genes with at least one zero expression level. To evaluate how high operon-/pathway-level SCS is, we conducted permutation test and test statistical significance. In permutation test, Combinations of genes in each operon/pathway were randomly shuffled to generate 10,000 false data sets. We calculated averaged operon-/pathway-level SCS defined as below

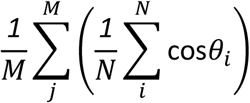

where 𝑀 is the number of operons/pathways. The value given in parentheses is operon-/pathway-level SCS. The empirical *p*-value was computed as <u>(𝑏 +</u> *<u>1</u>*<u>)/(𝑚 +</u> *<u>1</u>*) where 𝑏 is the number of permutations that gave the values equal to or lower than the original operon-/pathway-level SCS, and 𝑚 is the total number of permutations (10,000).

To evaluate stoichiometry conservation strength with the deleted gene, we excluded the three strains (Δ*bioA*, Δ*pabA*, and Δ*ygfZ*) with large expression changes caused by the deletion to ensure that these changes did not affect the calculation of stoichiometry conservation strength, and performed the calculation using the remaining 19 strains. Similarly, for the human data, the stoichiometry conservation strength between the differentially expressed genes (DEGs) and the perturbed gene of interest was calculated under the condition of excluding the perturbed gene, using 𝑁 − *1* genes from the total gene set (𝑁).

### DSC Score Analysis

To verify whether there is a statistically significant difference in the stoichiometry conservation strength with the perturbed gene between DEGs and non-DEGs, we compared the distribution of stoichiometry conservation strength between upDEGs/downDEGs and the perturbed gene, as well as between all genes and the perturbed gene, using the Brunner--Munzel test (*38*). This was performed in MATLAB with the “approxbrunnermunzel” function (v 2.0). If the 𝑝-value was below 0.05, the result was considered significant. The relative treatment effect 𝑃(𝑋 > 𝑌) − 𝑃(𝑋 < 𝑌), where 𝑋 (𝑌) is the set of DEGs (all genes) and 𝑃(𝑋 > 𝑌) is the probability that SCS of 𝑋 is larger than SCS of 𝑌, was defined as DSC score (DEG Stoichiometry Conservation score). DSC score is calculated for both upDEGs and downDEGs under a certain perturbation condition and is defined as uDSC score and dDSC score, respectively. When DSC score was greater than 0, it indicated that the DEGs significantly conserved their stoichiometry with the perturbed gene, and when it was lower, it indicated that they did not conserve it, because 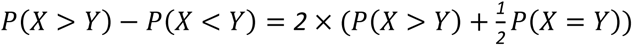.

### Generality of DSC Score Analysis

Genome-Wide Perturb-Seq data were downloaded from (*13*). We used K562 genome-scale perturb-seq data of the three datasets. Stoichiometry conservation network was constructed by using transcriptome data with large transcriptional change (≥ *200* DEGs) excluding the perturbation data of interest. DEGs were defined as genes with significance of 𝑝 < *0*.*05* by Anderson-Darling test following Benjamini-Hochberg correction. Worm Perturb-Seq data were downloaded from Gene Expression Omnibus under the accession number GSE253847.

Stoichiometry conservation network was constructed by using transcriptome data with large transcriptional change (≥ *5* DEGs) excluding the perturbation data of interest. DEGs were defined as genes with significance of FDR < *0*.*05* by EmpirDE (*40*).

