## Supplementary figures for "Global propagation of genetic perturbation effects through genome-wide stoichiometry conservation architecture"

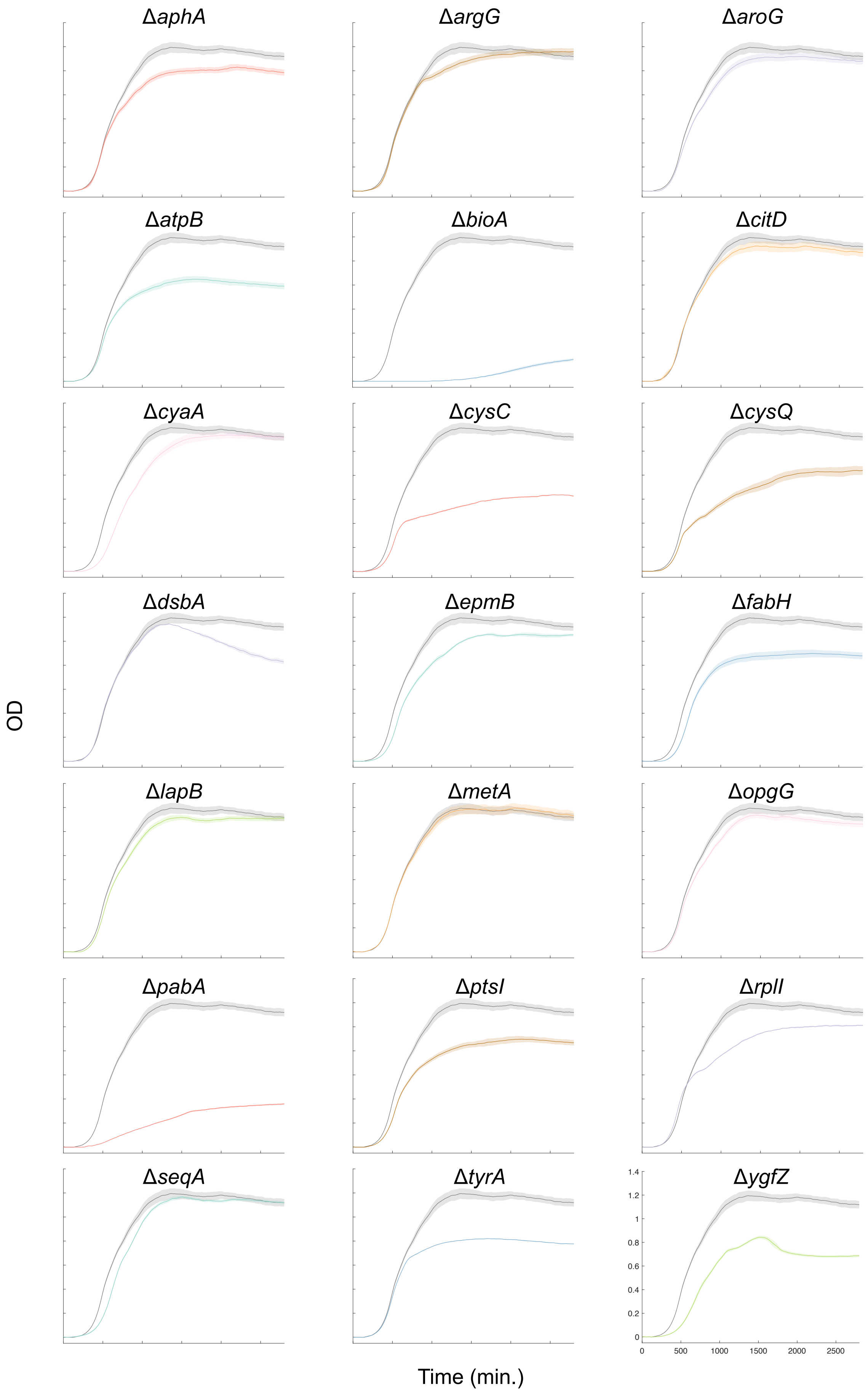


**Supplementary figure S1. Growth curves.**

Population growth curve of parental strain BW25113 (grey) and the single-gene deletion strains. Standard errors are shown as shading above and below the mean.

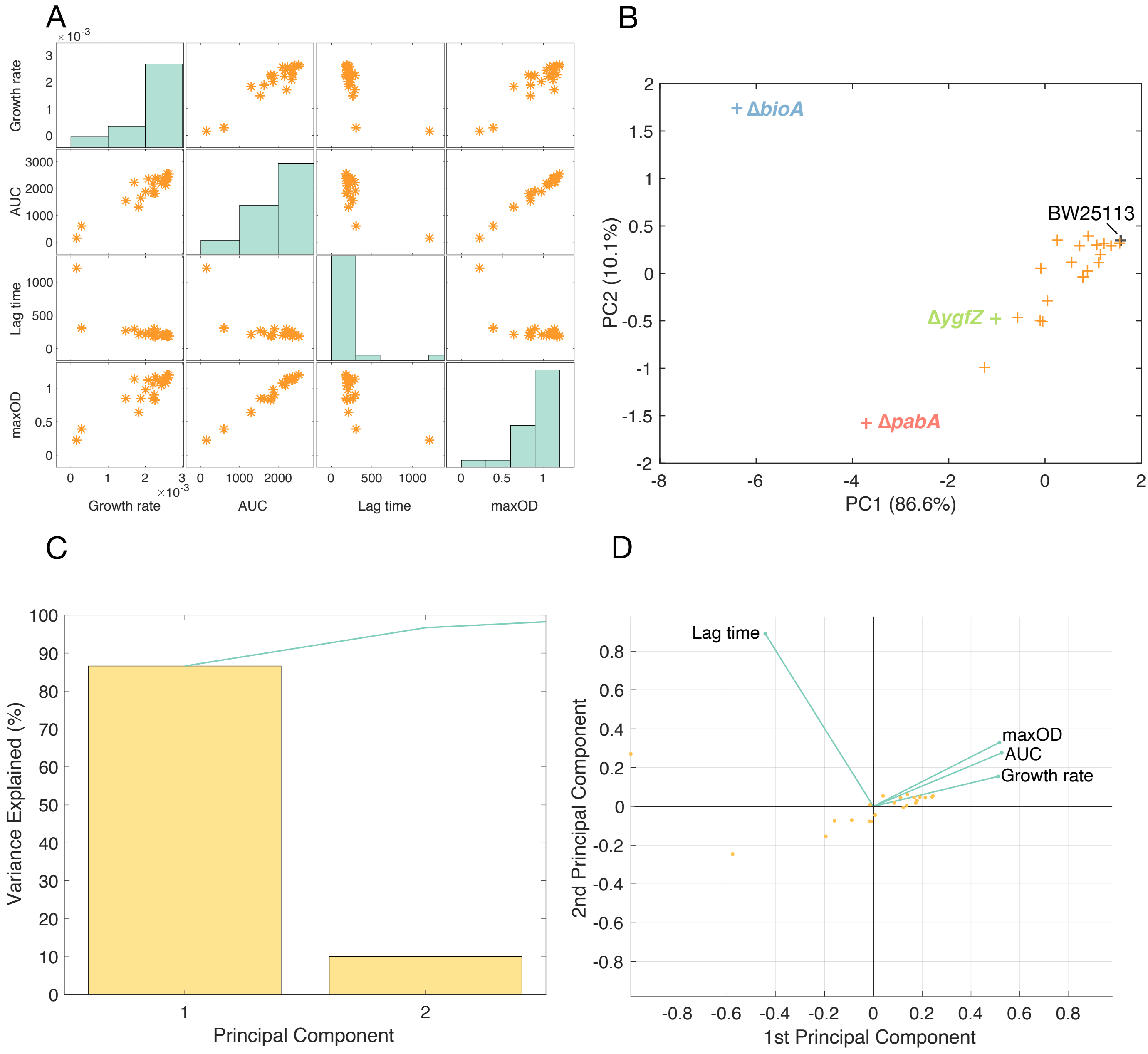


**Supplementary figure S2. PCA analysis of growth parameters.**

(**A**). Matrix plot of four growth parameters. The subplot in the $i$-th row, $j$-th column is a scatter plot of the $i$-th parameter and $j$-th parameter. Each dot corresponds to one strain. Along the diagonal are histogram plots of each growth parameter. (**B**). Scatter plot of first two principal component. (**C**). The percentage of variance explained by each principal component. The first two components explain approximately 95% of the variance of the data. (**D**) A orthonormal principal component coefficients of each variable and principal component scire of each observation shown in the PCA space.

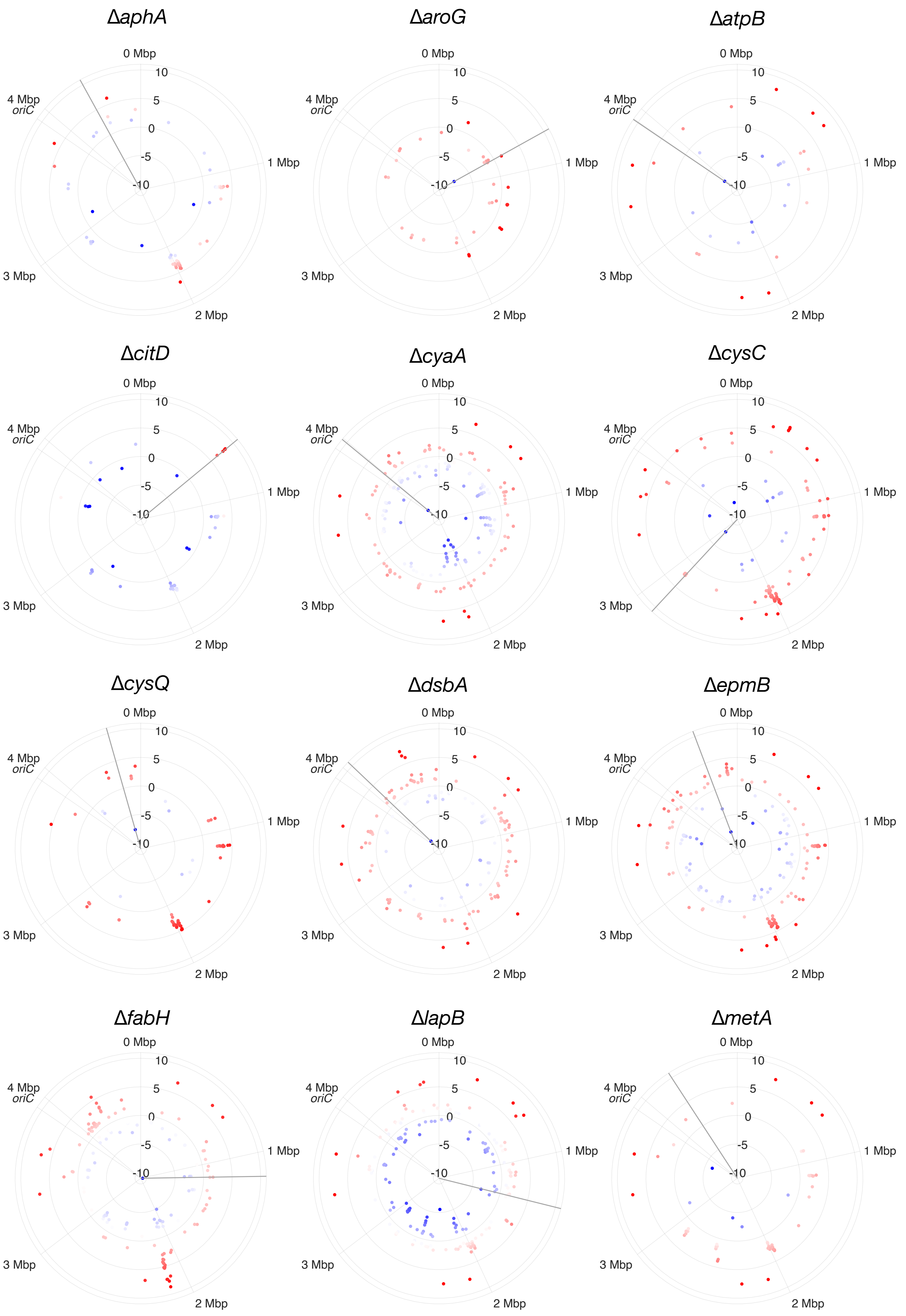
ΔaphA
ΔaroG
ΔatpB
ΔcitD
ΔcyaA
ΔcysC
ΔcysQ
ΔdsbA
ΔepmB
ΔfabH
ΔlapB
ΔmetA
0 Mbp
1 Mbp
2 Mbp
3 Mbp
4 Mbp
oriC
10
5
0
-5
-10

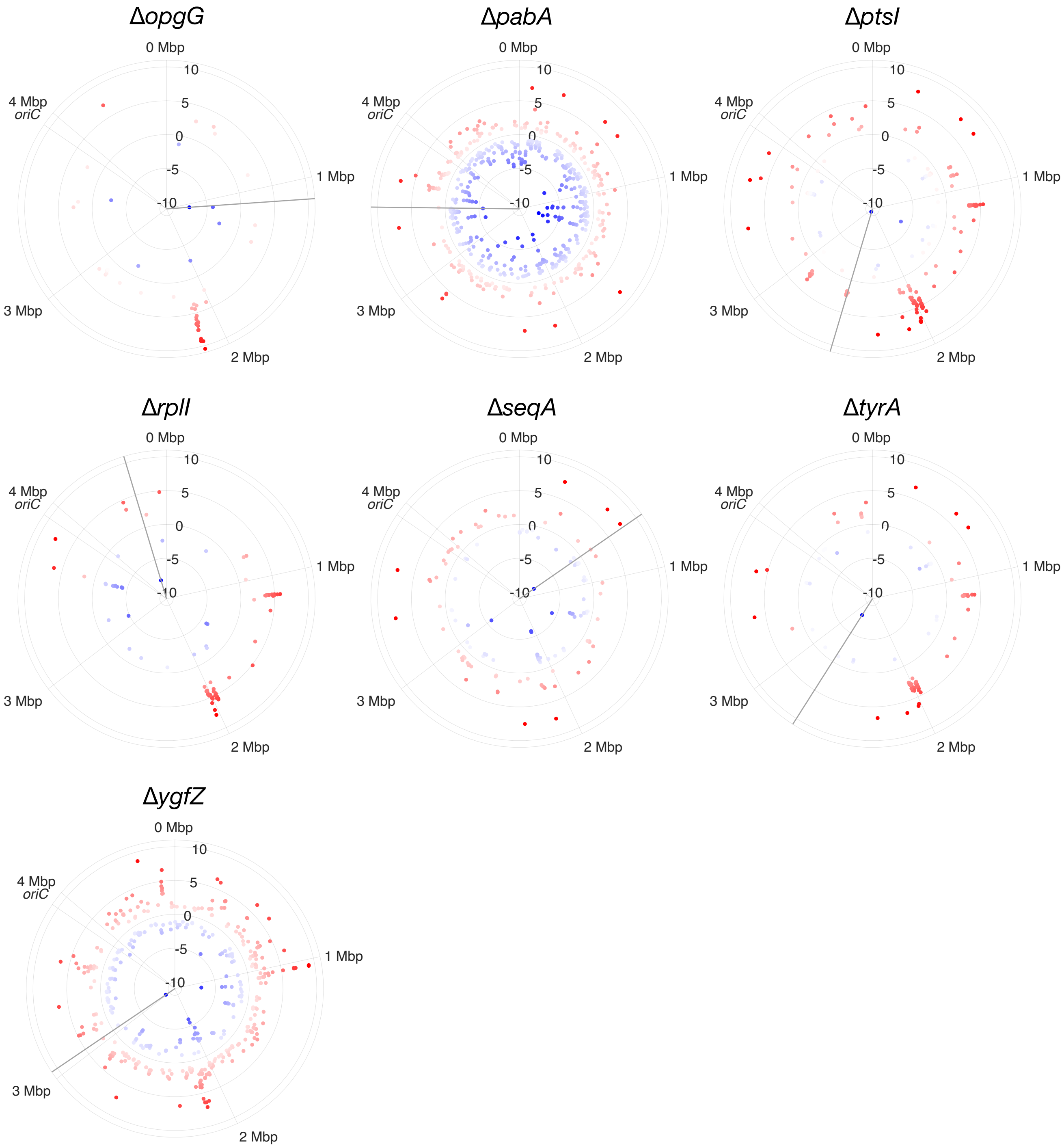


**Supplementary figure S3. DEGs mapping to *E. coli* chromosome**

Differentially expressed genes (DEGs) detected in each deletion strain are mapped to the locus in *E. coli* chromosome. upDEGs and downDEGs are represented by red/blue color respectively. Polar radius represents the chromosomal location, and radius distance represents log fold change of expression level relative to the BW25113 strain.

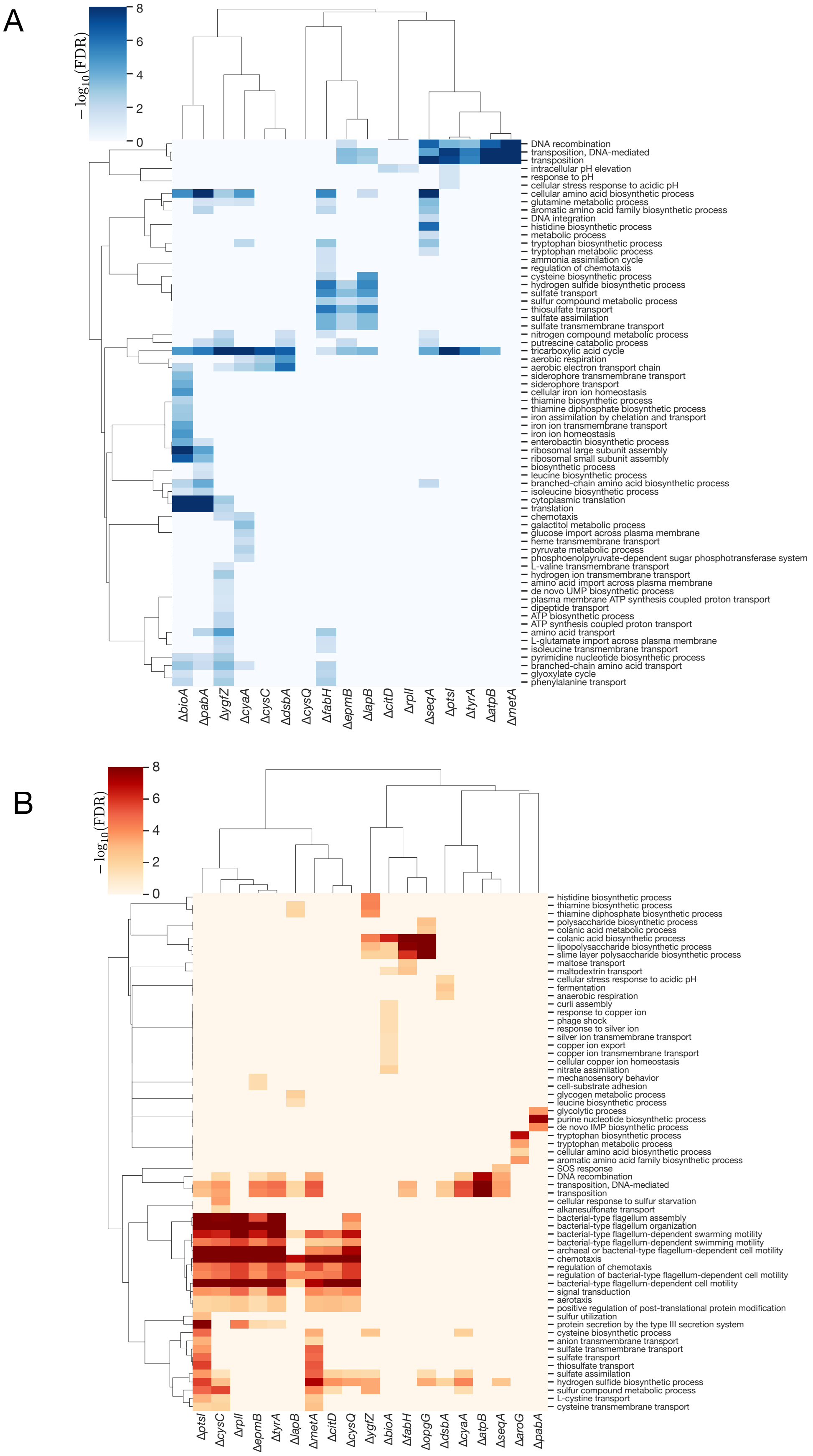


**Supplementary figure S4. GO enrichment analysis and hierarchical clustering.**

GO terms enriched in each deleted strains were hierarchical clustered based on its FDR value. Column labels correspond to the strains, and row labels correspond to the enriched GO terms.

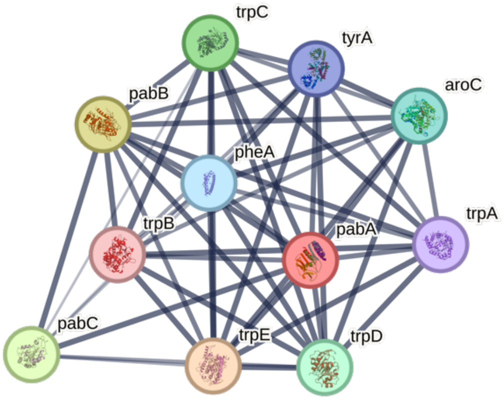


**Supplementary figure S5. Genes encoding proteins that functionally or physically interact with *pabA*.** Edges between genes indicate physical or functional interactions between two proteins, and the thickness of the edge indicates the strength of data support for the interaction between the two proteins. Figure taken from STRING (59).

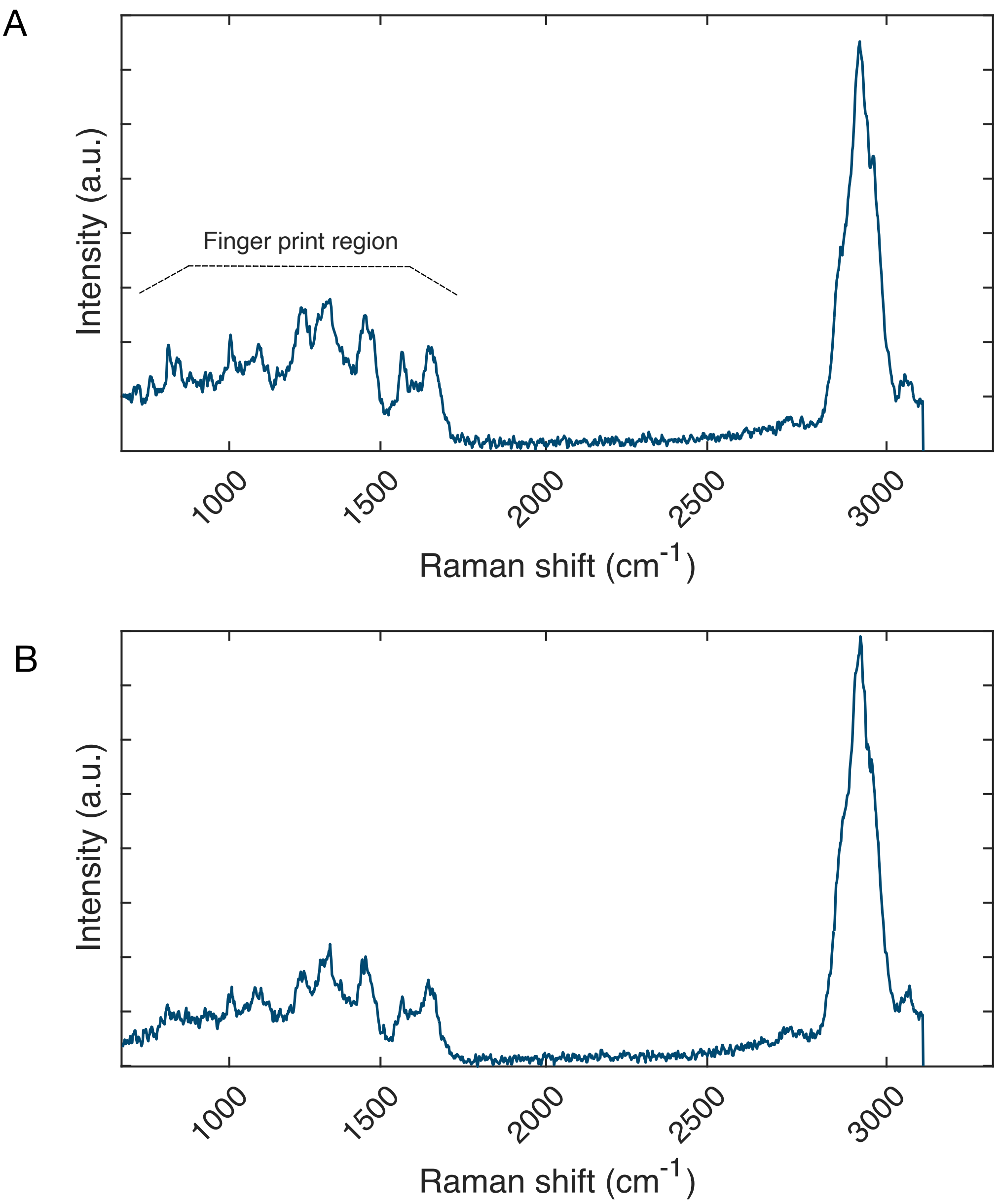


**Supplementary figure S6. Representative single-cell Raman spectra.** Single-cell level Raman spectra obtained from (A) BW25113 and (B) Δ*bioA* strain.

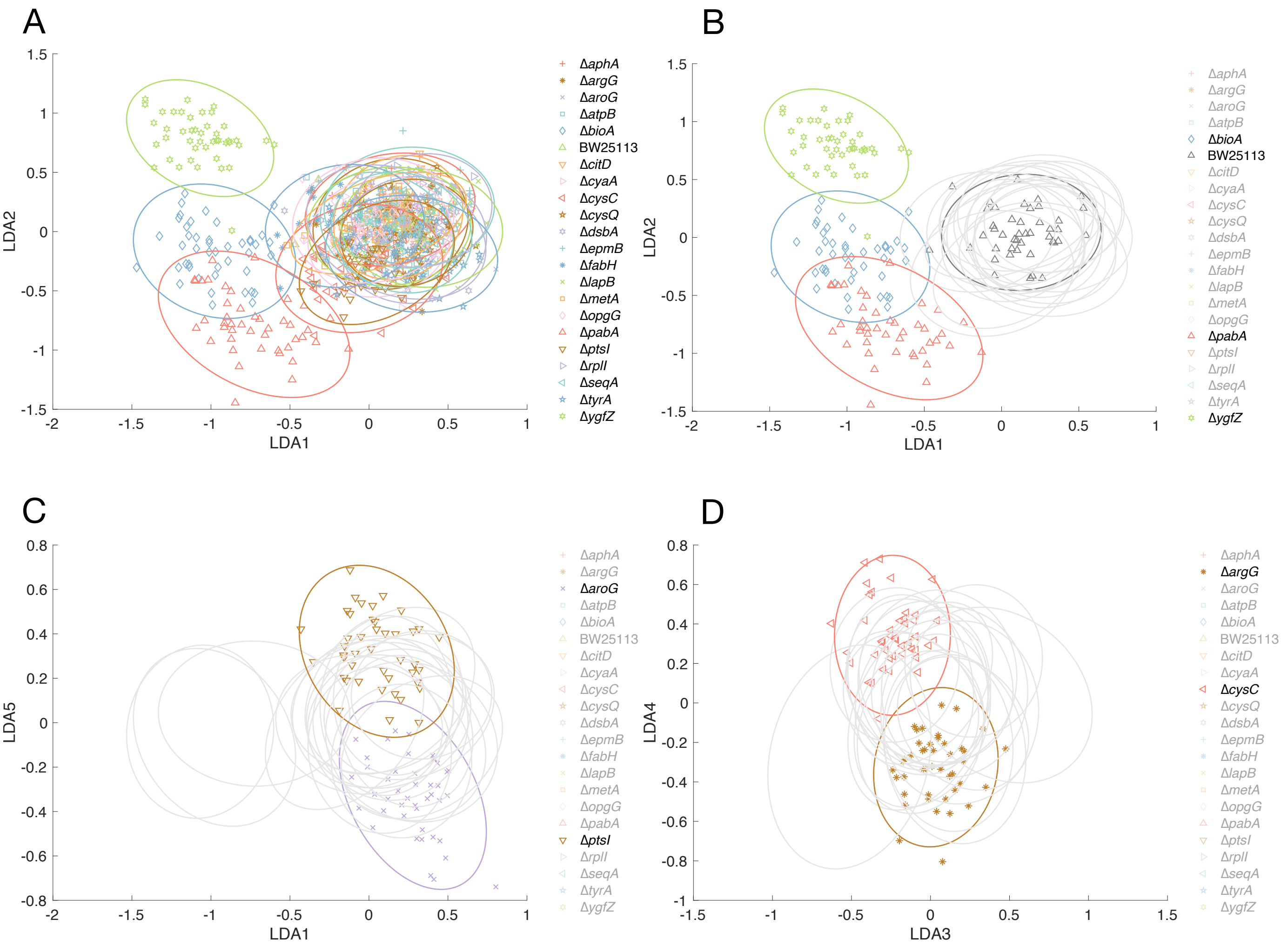


**Supplementary figure S7. Raman spectral analysis of *E. coli* single gene deletion strains.** (**A-D**). Cellular Raman spectra in LDA space. The dimensionality of the spectra is reduced to 21 (=22-1). Each data point represent spectrum from a single cell, and each ellipse shows the 95% concentration ellipse for each condition. Their projection to the LDA1-LDA2 plane (A, and B), the LDA1-LDA5 plane (C), and the LDA3-4 plane (D) are shown. Δ*bioA*, Δ*pabA*, Δ*ygfZ* formed distinct clusters separated from other strains. Δ*aroG* and Δ*ptsI* (C), and Δ*argG* and Δ*cysC* (D) are distinguishable each other.

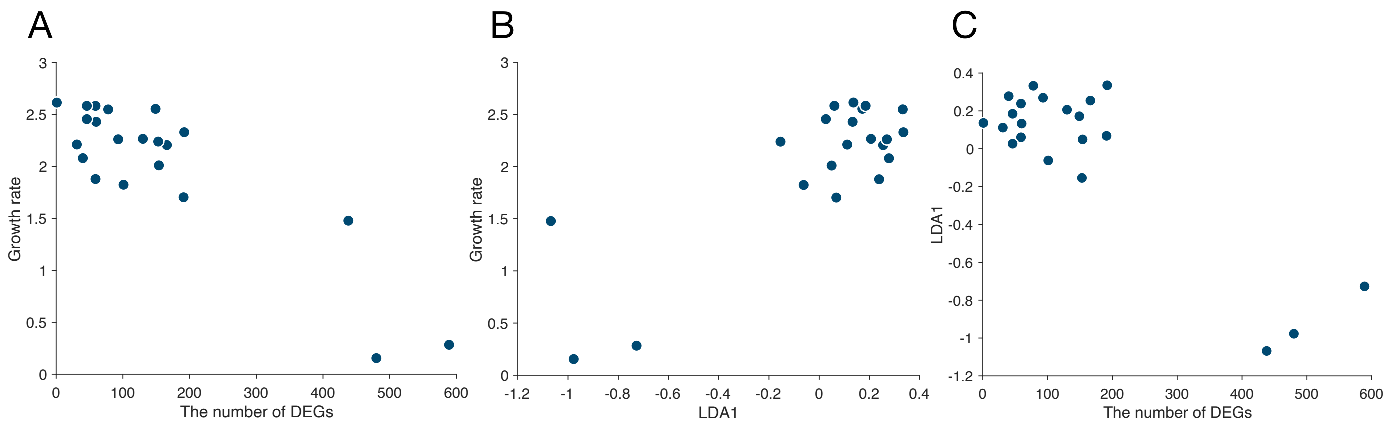


**Supplementary figure S8. Growth, transcriptome profile, and Raman spectral changes caused by single-gene deletions.** (**A-C**). Scatter plot between growth rate and the number of DEGs (A), growth rate and the value of LDA1 axis (B), and the value of LDA1 axis and the number of DEGs (C).

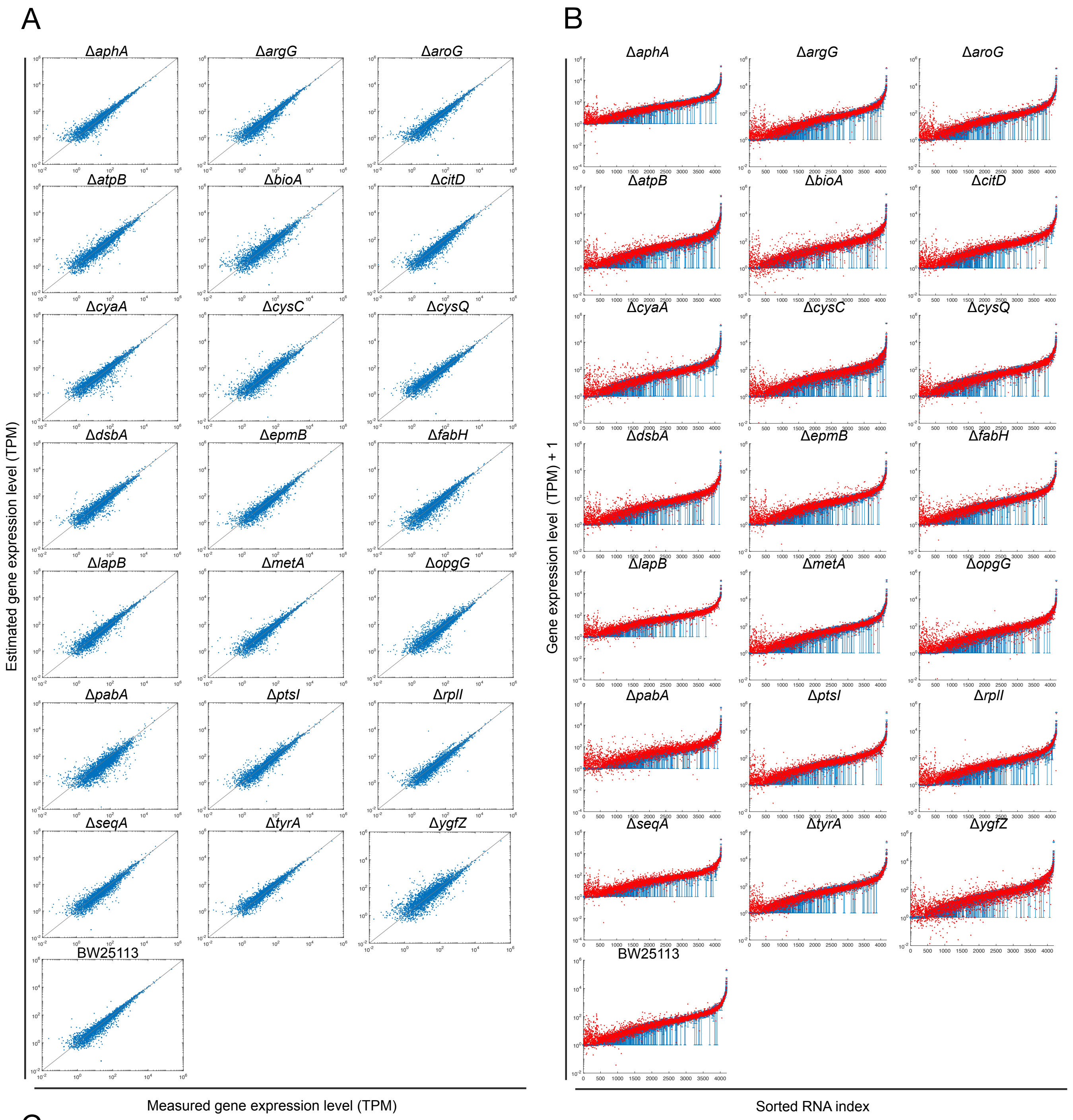


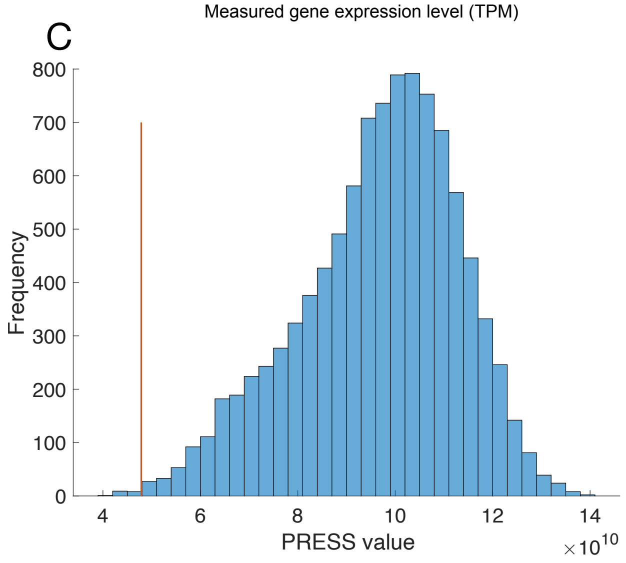


**Supplementary figure S9.**

**Estimation of transcriptomes from Raman spectra.**

(**A and B**). Estimation of transcriptomes from Raman spectra in the LDA space. (A) is scatter plots between measured and estimated gene expression level (TPM). Straight lines represent $y = x$. In (B), blue points represent the measured gene expression level (average of the replicate measurements; error bar, SE), which are sorted along the horizontal axis. Red points represent gene expression level estimated from Raman spectra. (**C**) Histogram of $PRESS_t$ values of randomly permuted data. The $PRESS_t$ of the original dataset is $4.79 \times 10^{10}$. The p-value was 0.0020.

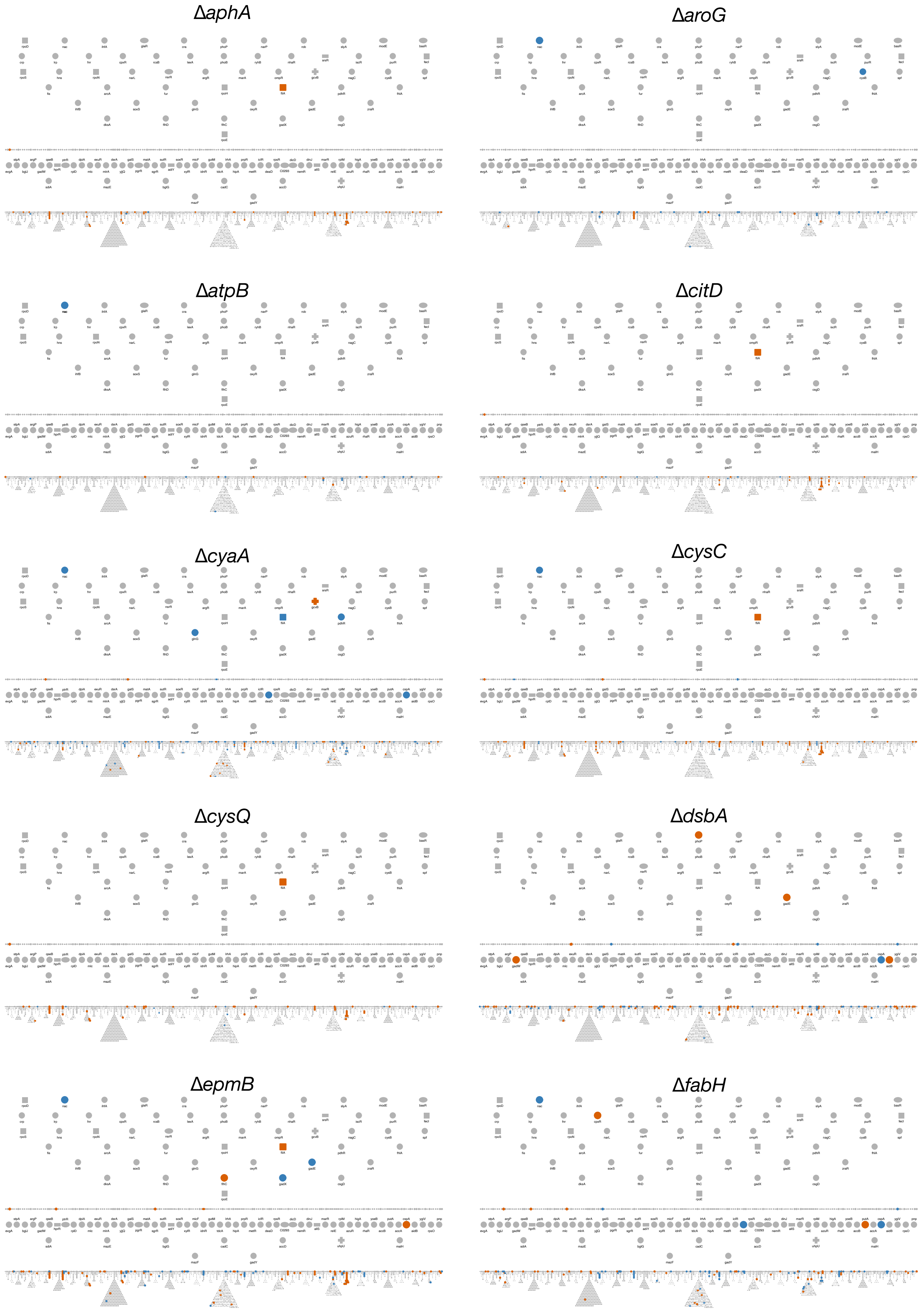
ΔaphA
ΔaroG
ΔatpB
ΔcitD
ΔcyaA
ΔcysC
ΔcysQ
ΔdsbA
ΔepmB
ΔfabH

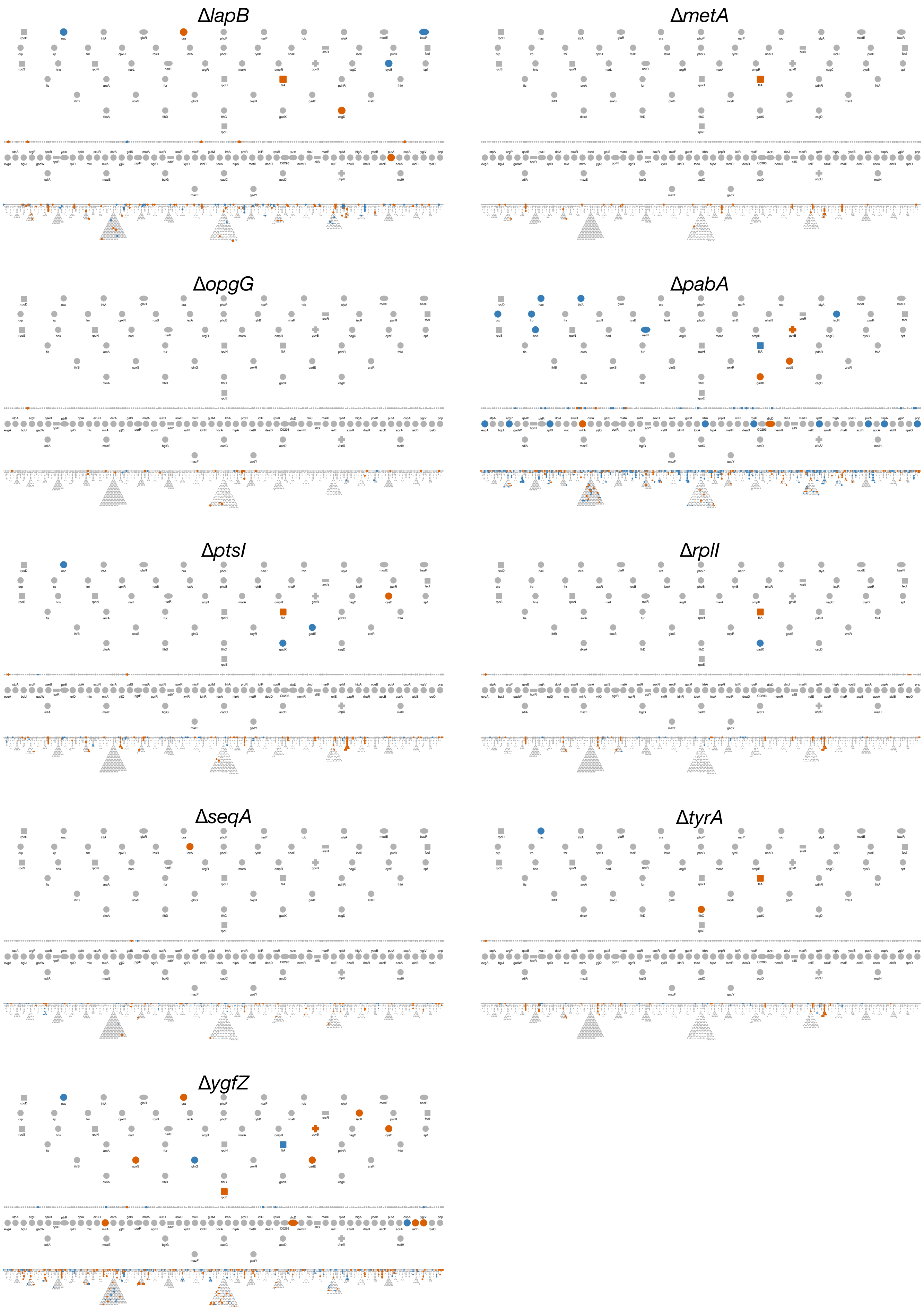


**Supplementary figure S10. DEGs mapping to gene regulatory network**

Differentially expressed genes (DEGs) detected in each deletion strain are mapped to gene regulatory network obtained from EcoCyc. upDEGs and downDEGs are represented by red/blue color respectively.

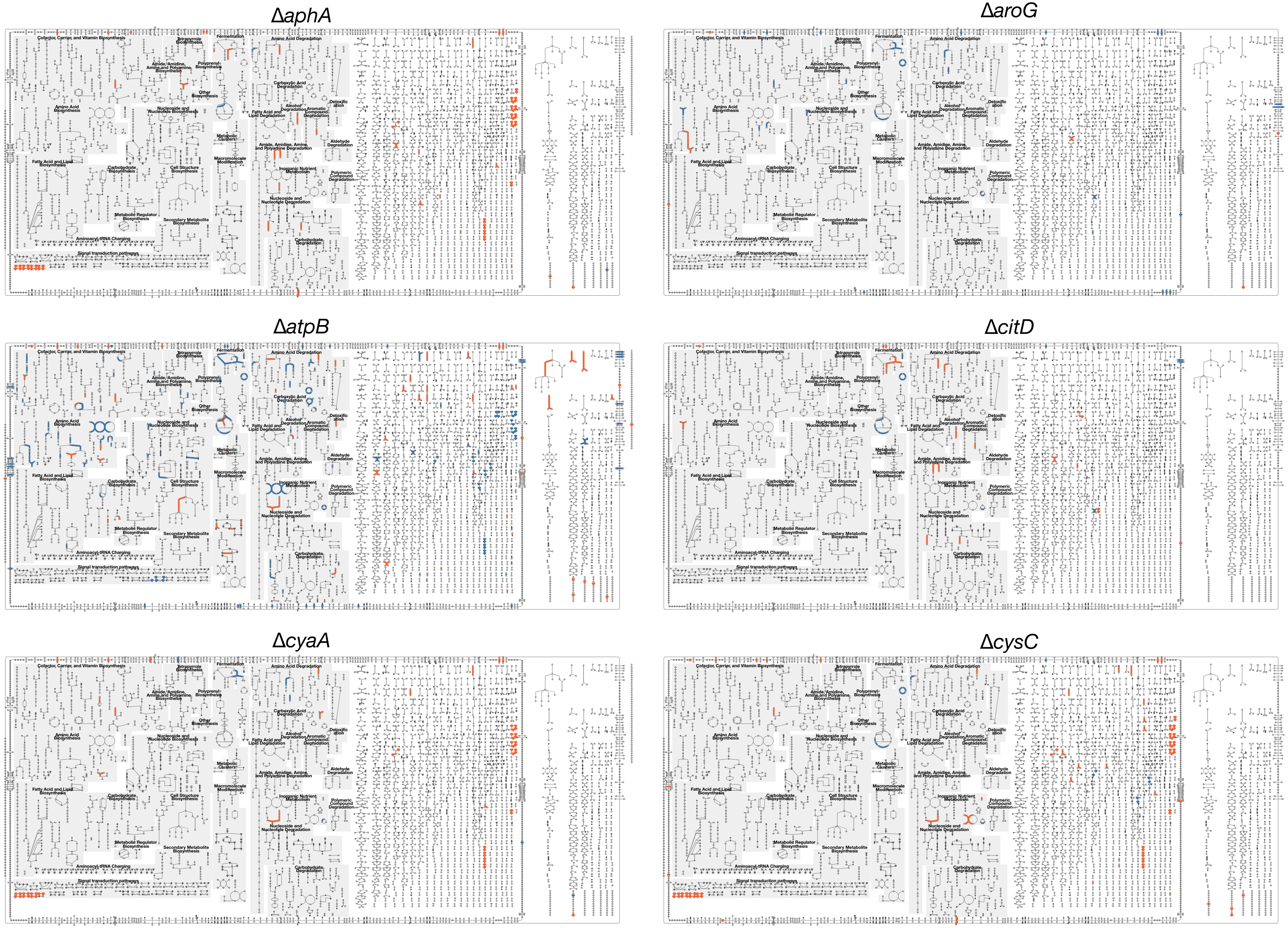
ΔaphA
ΔaroG
ΔatpB
ΔcitD
ΔcyaA
ΔcysC

### Δ*cysQ*

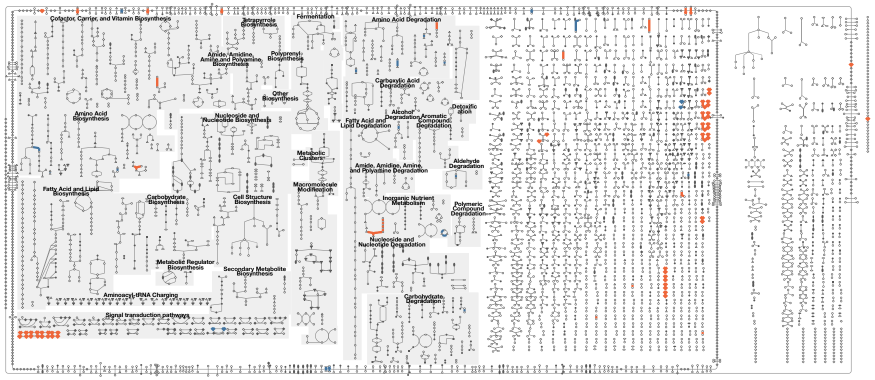

### Δ*dsbA*

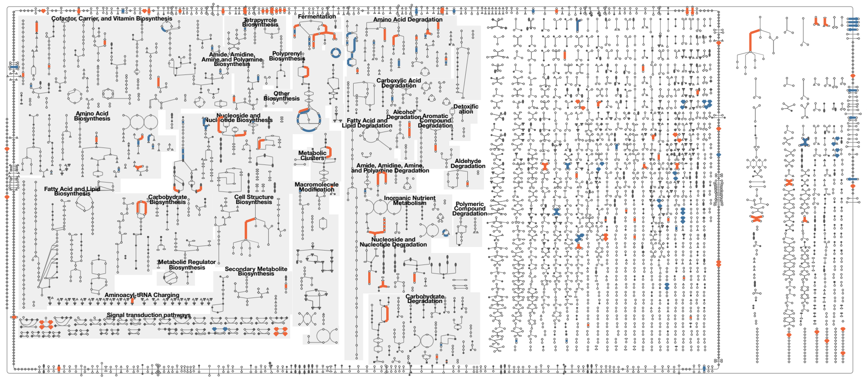

### Δ*epmB*

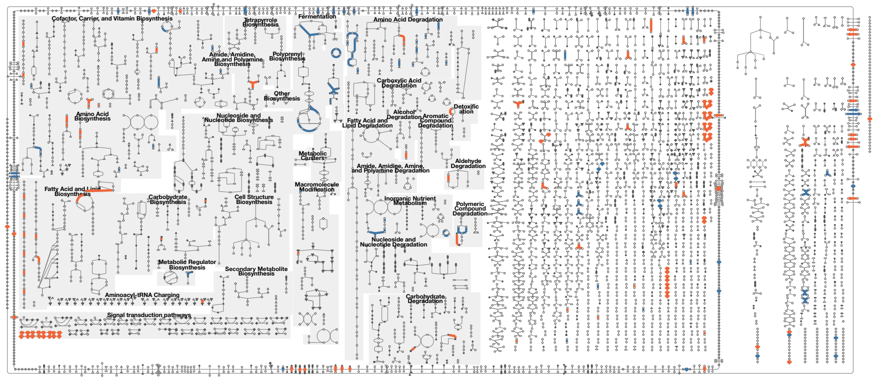

### Δ*fabH*

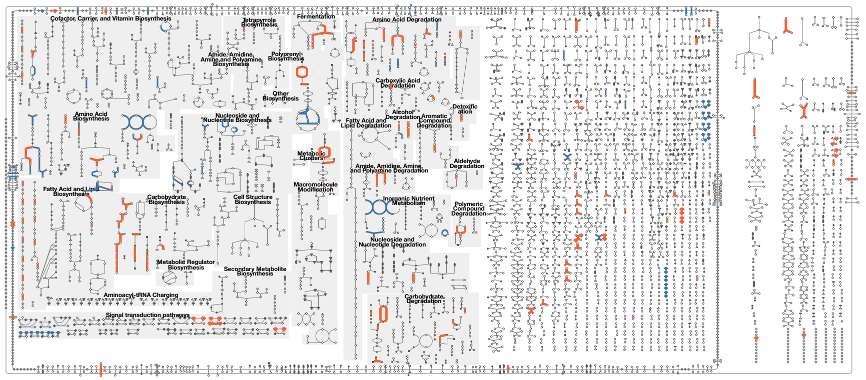

### Δ*lapB*

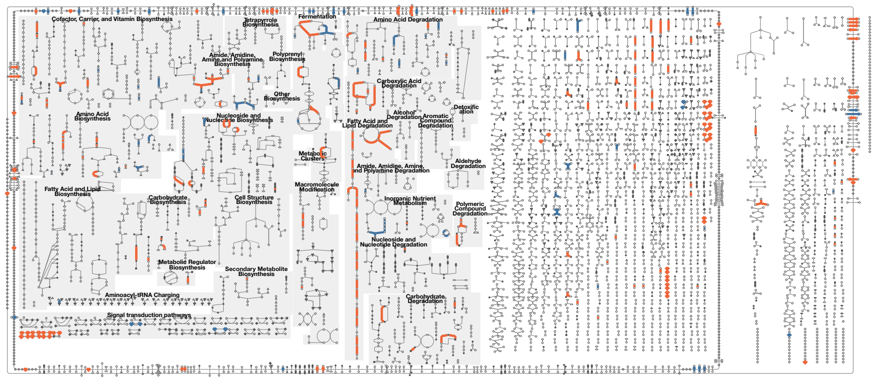

### Δ*metA*

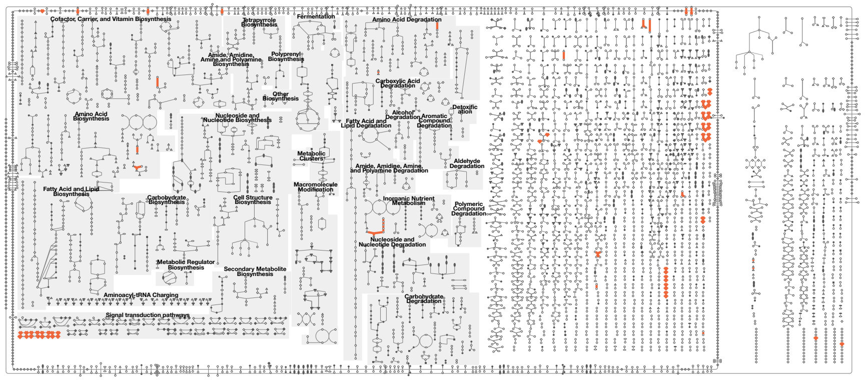

*ΔopgG*

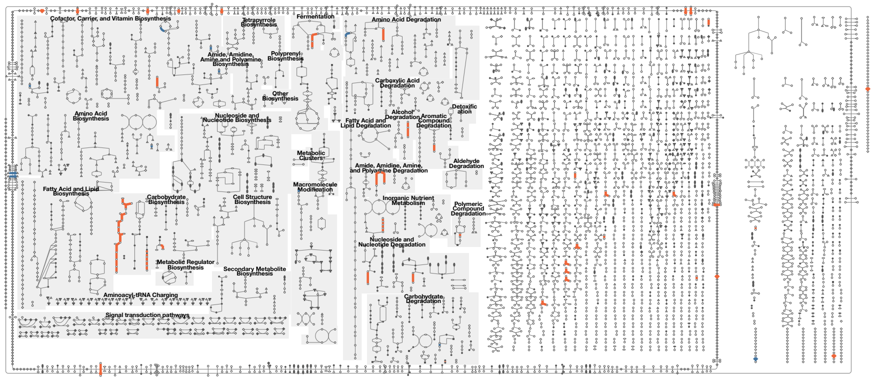

*ΔpabA*

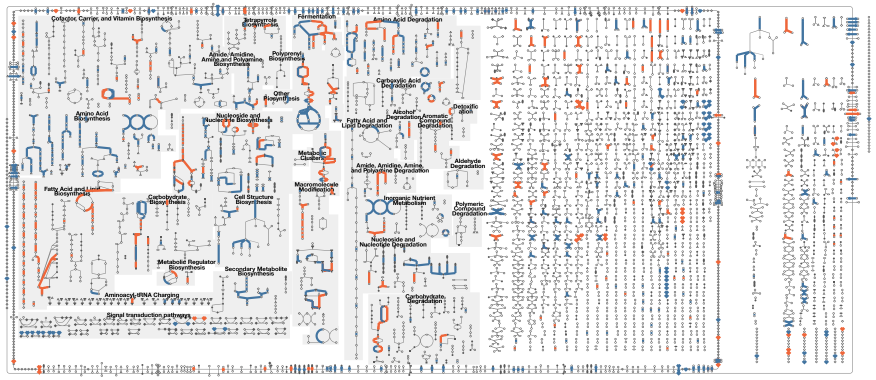

*ΔptsI*

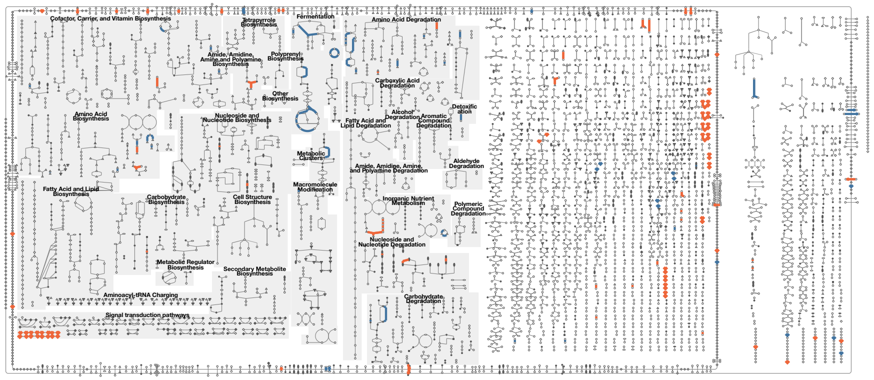

*ΔrplI*

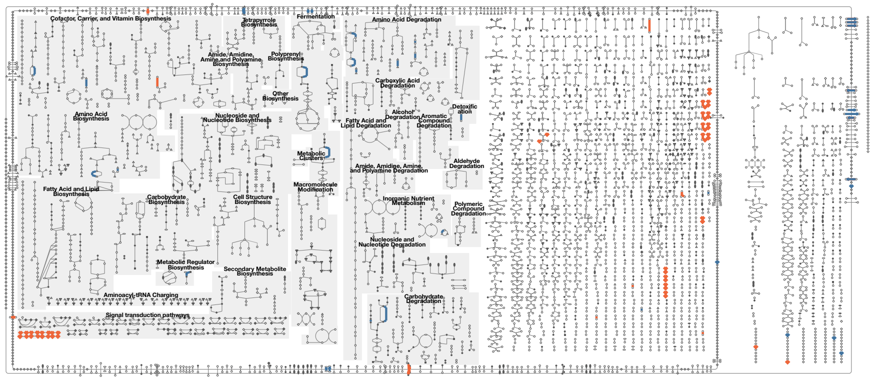

*ΔseqA*

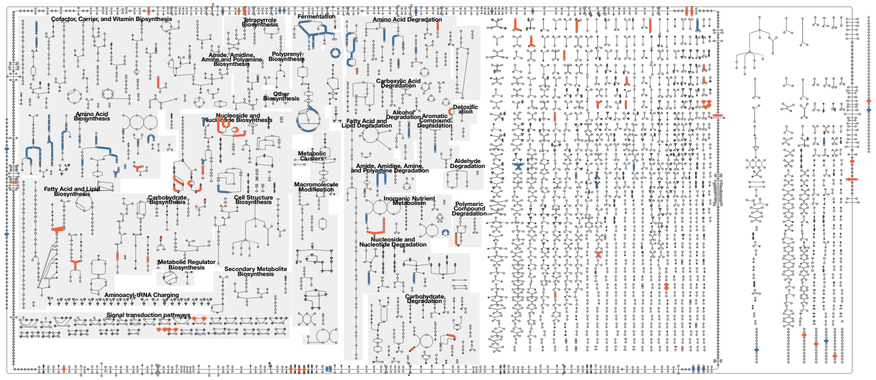

*ΔtyrA*

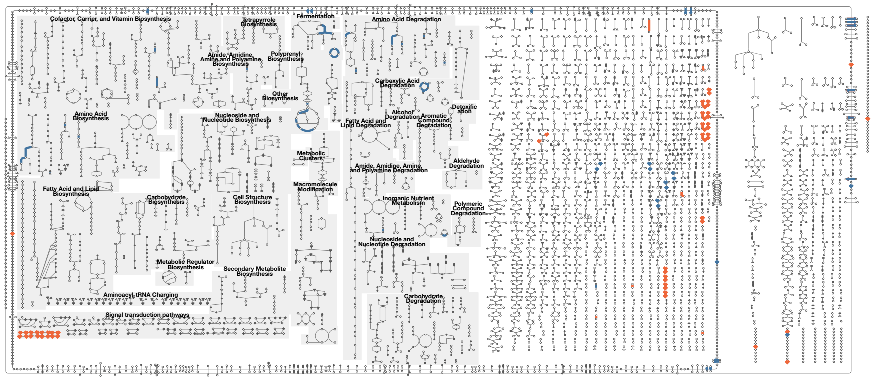

Δ*ygfZ*

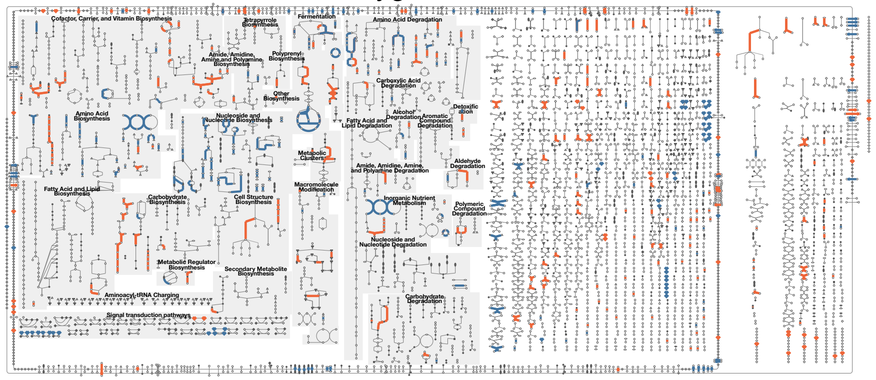

**Supplementary figure S11. DEGs mapping to metabolic network.**

Differentially expressed genes (DEGs) detected in each deletion strain are mapped to metabolic network obtained from EcoCyc. upDEGs and downDEGs are represented by red/blue color respectively.

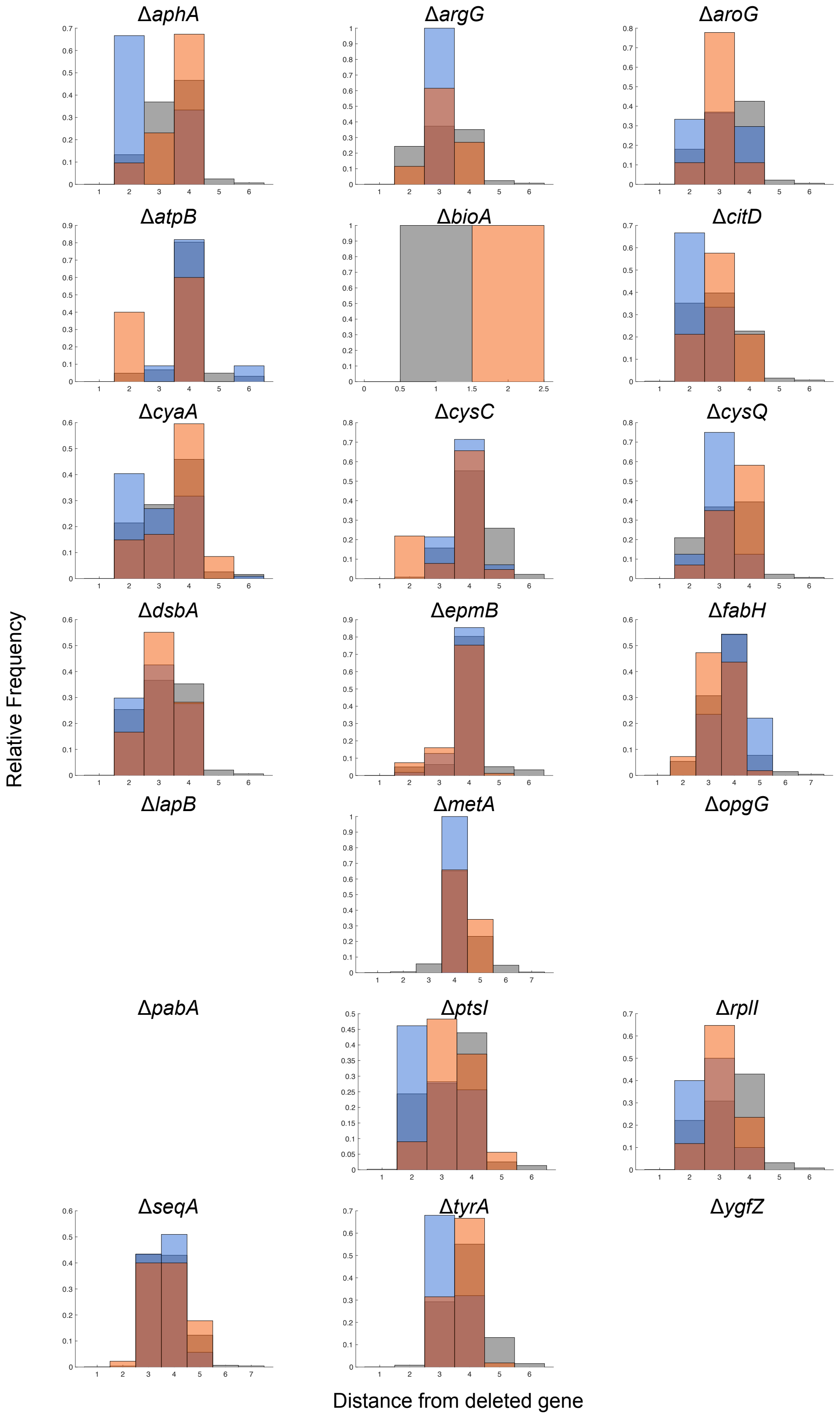


**Supplementary figure S12. Histogram of network distances between the deleted gene and DEGs in gene regulatory network.** The distance between deleted genes and upDEGs (red), downDEGs (blue) and all genes (gray) in each deletion strain is displayed as a histogram. Δ*lapB*, Δ*opgG*, Δ*pabA*, and Δ*ygfZ* are left blank because they were not included in the original network data.

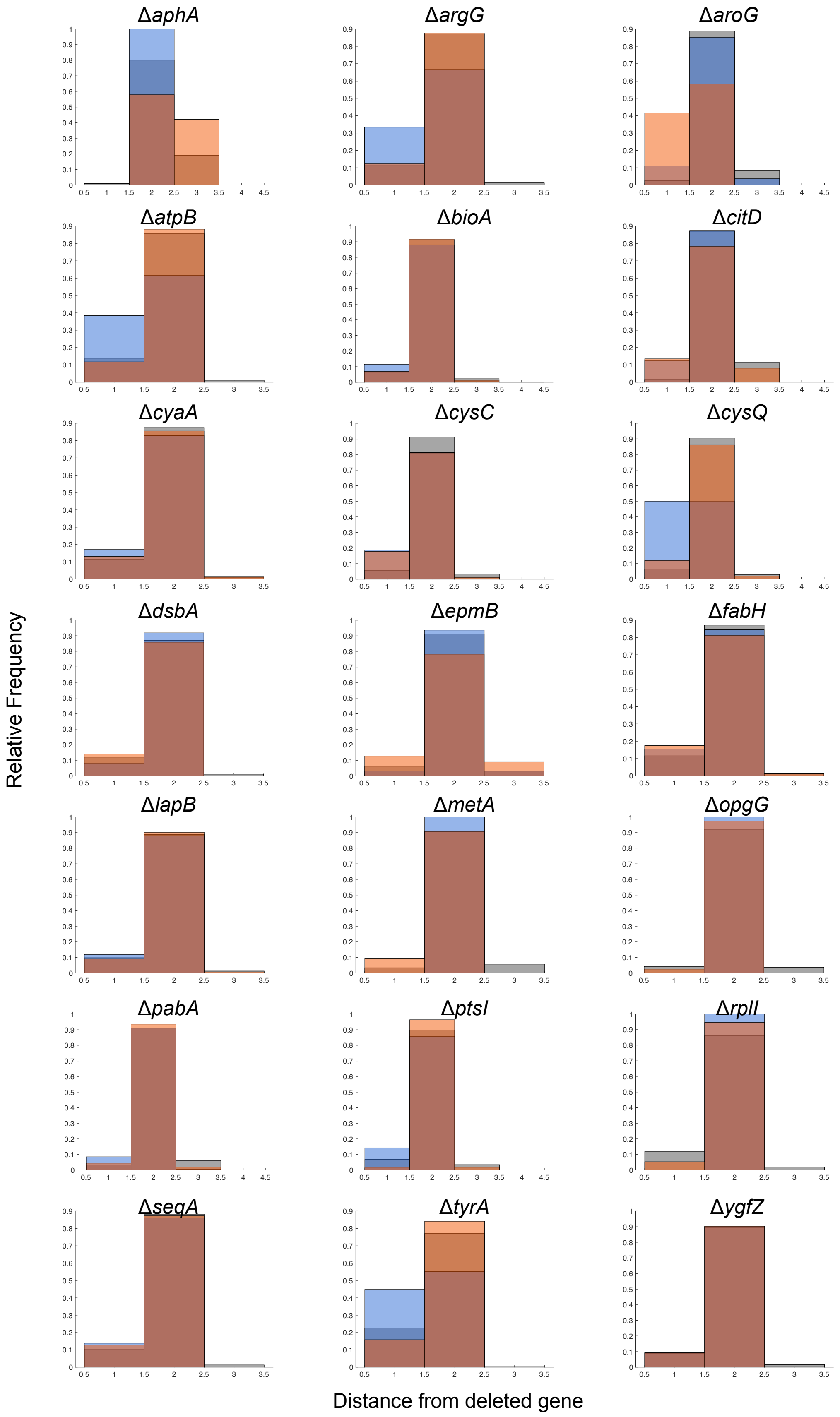


**Supplementary figure S13. Histogram of network distances between the deleted gene and DEGs in functional protein interaction network.** The distance between deleted genes and upDEGs (red), downDEGs (blue) and all genes (gray) in each deletion strain is displayed as a histogram.

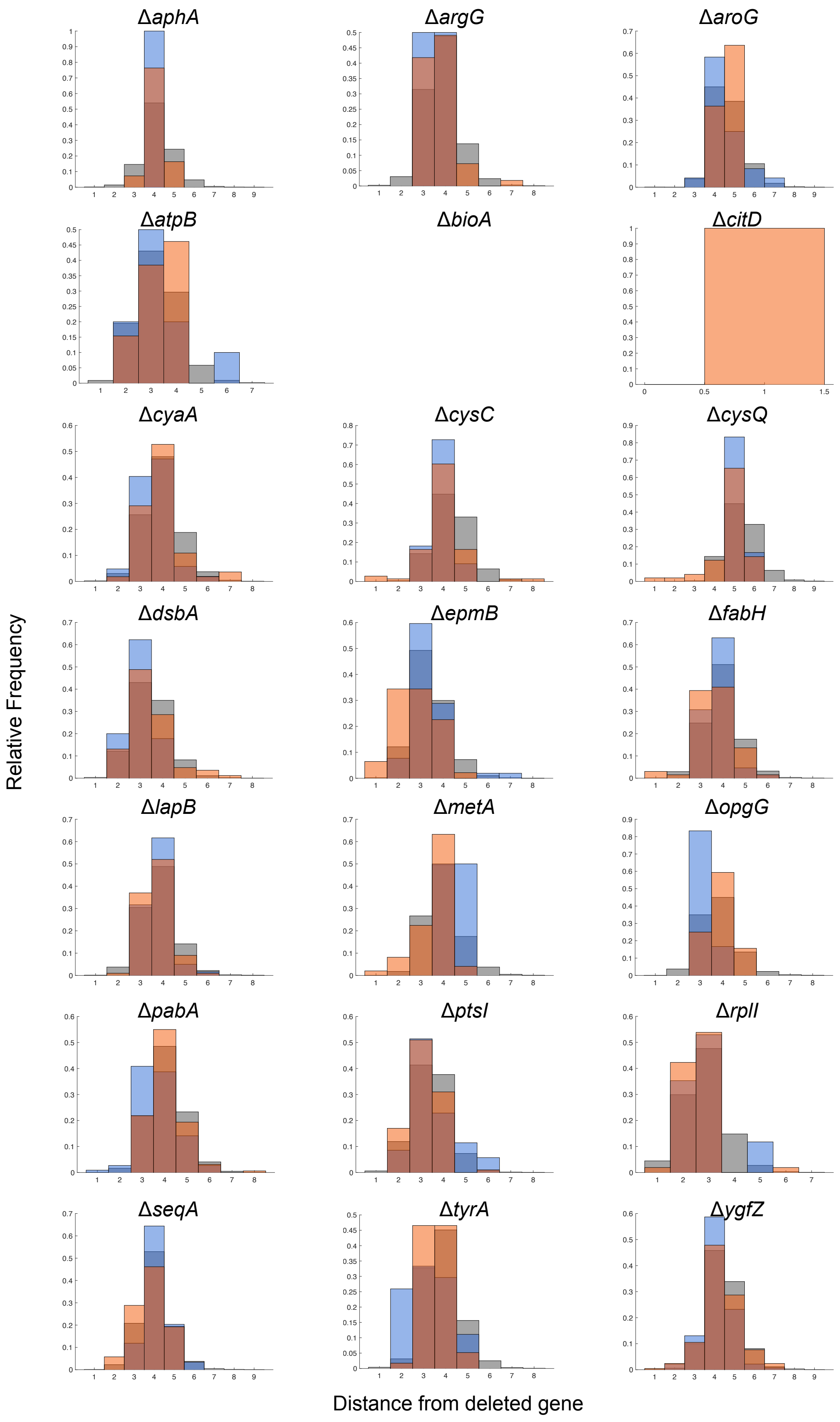


**Supplementary figure S14. Histogram of network distances between the deleted gene and DEGs in physical protein interaction network.** The distance between deleted genes and upDEGs (red), downDEGs (blue) and all genes (gray) in each deletion strain is displayed as a histogram. Δ*bioA* are left blank because they were not included in the original network data.

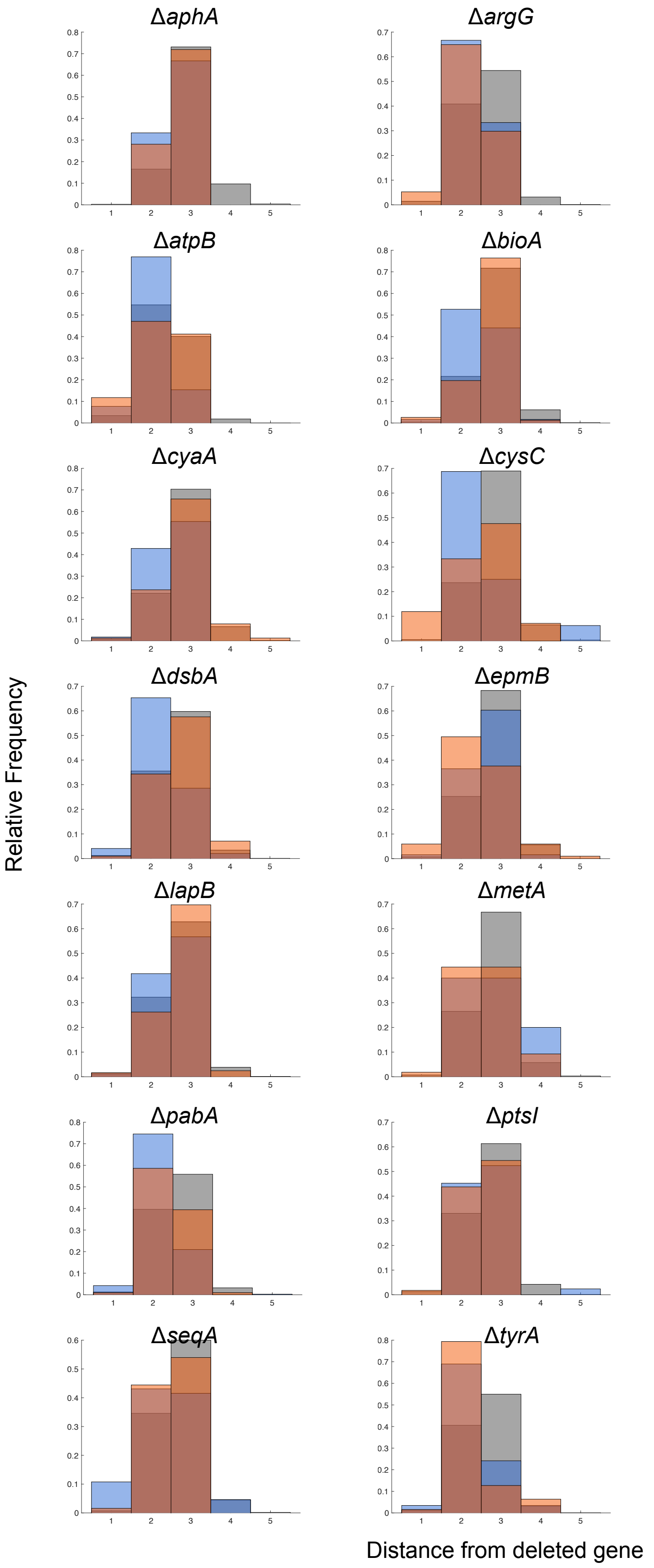


**Supplementary figure S15. Histogram of network distances between the deleted gene and DEGs in all gene dunctional network.** The distance between deleted genes and upDEGs (red), downDEGs (blue) and all genes (gray) in each deletion strain is displayed as a histogram.

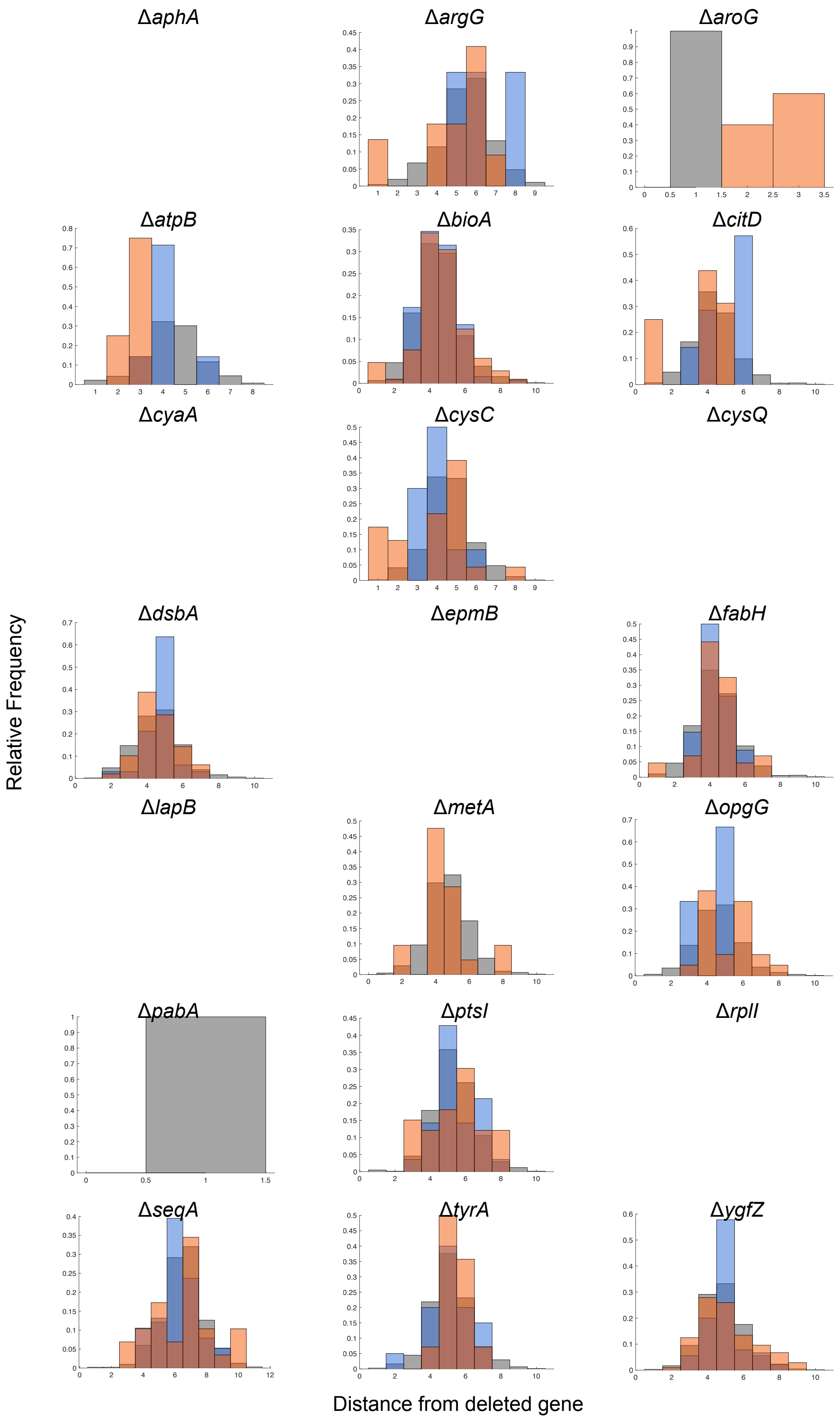


**Supplementary figure S16. Histogram of network distances between the deleted gene and DEGs in benchmark gene functional network.** The distance between deleted genes and upDEGs (red), downDEGs (blue) and all genes (gray) in each deletion strain is displayed as a histogram. *ΔaphA, ΔcyaA, ΔcysQ, ΔepmB, ΔlapB,* and *ΔrplI* are left blank because they were not included in the original network data.

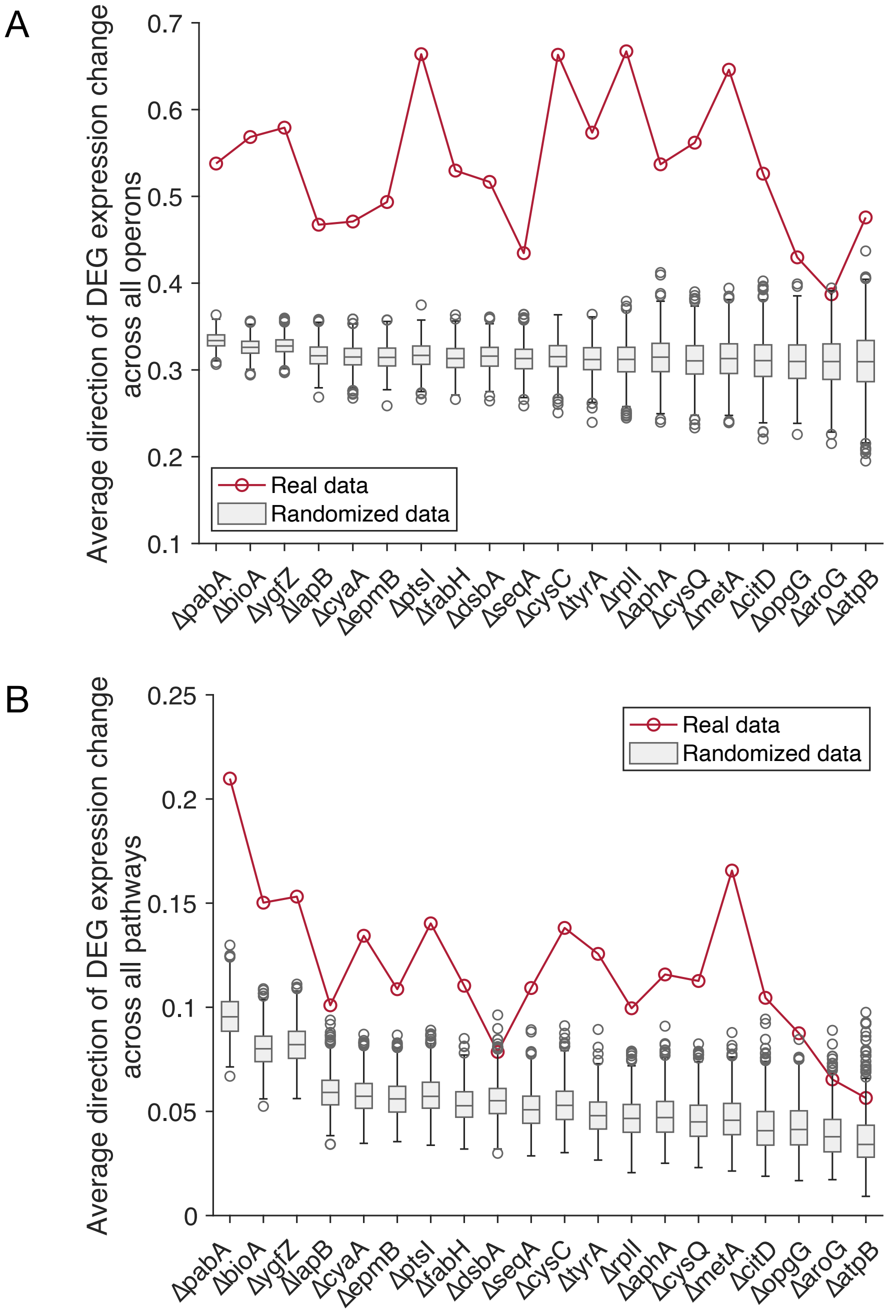


**Supplementary figure S17. Direction of DEG expression change.**

Average direction of DEG expression changes across operons (A) and pathway (B) was compared between real data (red) and randomized data (grey). 1,000 randomized data were generated. Empirical *p*-value is <0.001 for all strains in operon. Empirical *p*-value is <0.05 for almost all strains in pathway expect for Δ*atpB* (p=0.0729).

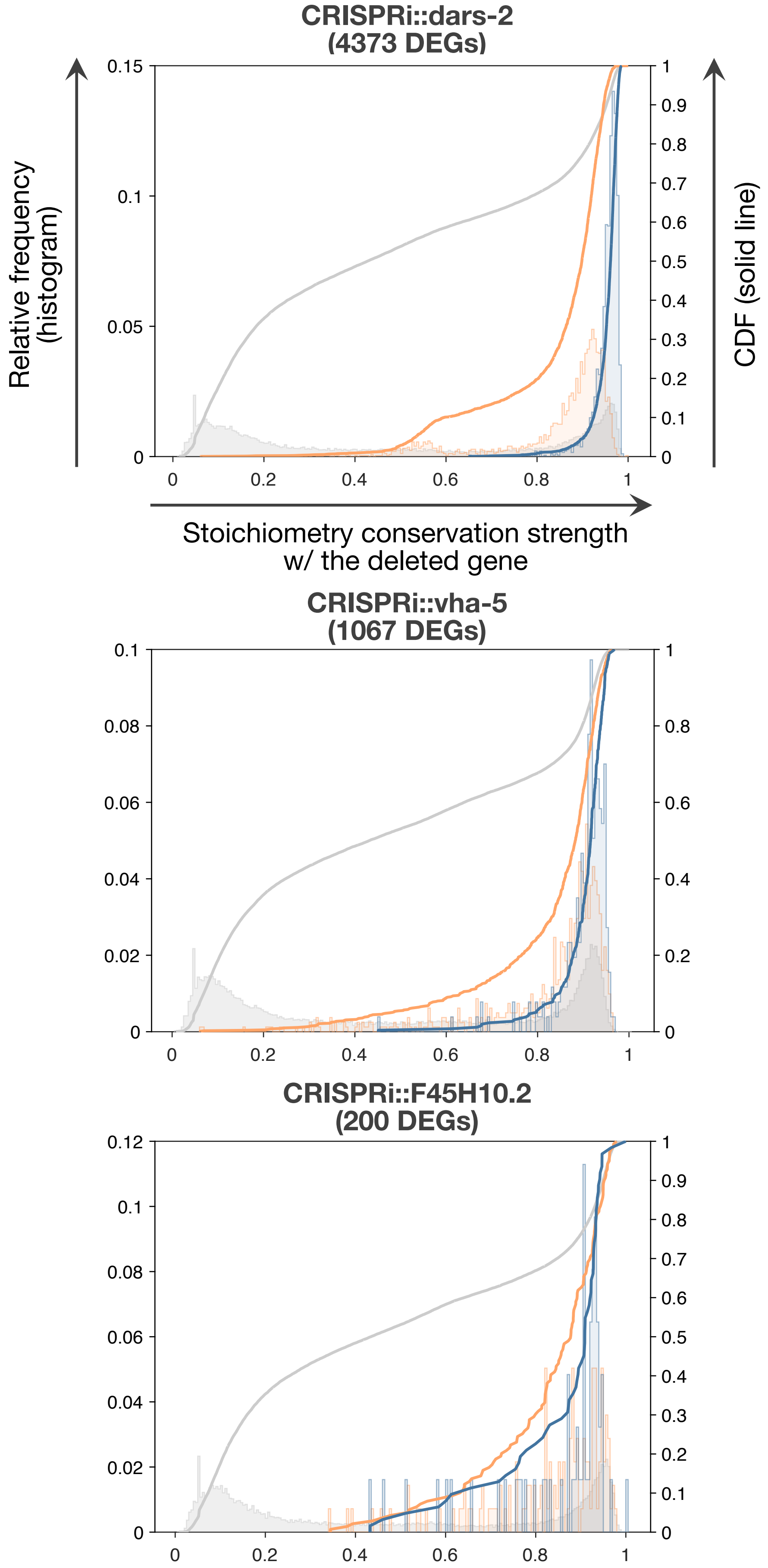


**Supplementary figure S18. Representative example of propagation of genetic perturbation effects through high SCS relation in *C. elegans* cells.** Same analysis in Fig. 5B and Fig. 6A.

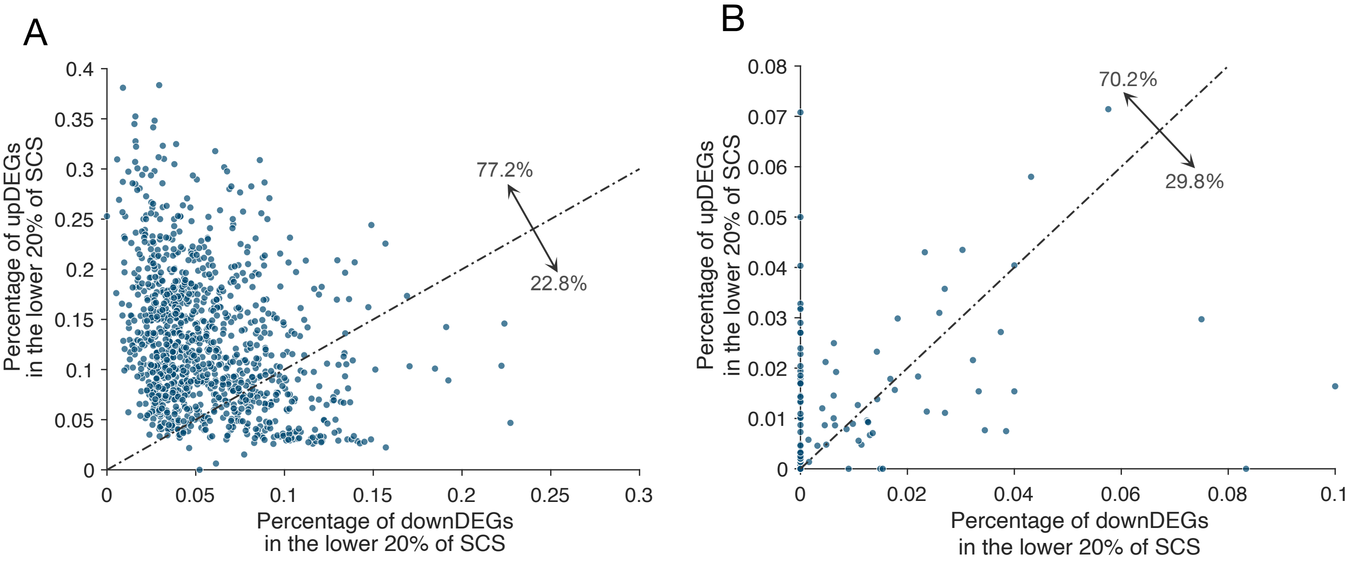


**Supplementary figure S19. Scatter plot of percentage of upDEGs and downDEGs in the lower 20% of SCS with the perturbed gene.** (**A**) human cell and (**B**) *C. elegans* cells. Each dot represents one perturbation. Dash-dotted line correspond to y=x. The percentage at the arrow represents the proportion of perturbation conditions above or below the y=x line. Points at (0,0) are excluded when calculating the proportions.
